# Evaluation of methods for estimating coalescence times using ancestral recombination graphs

**DOI:** 10.1101/2021.11.15.468686

**Authors:** Débora Y. C. Brandt, Xinzhu Wei, Yun Deng, Andrew H. Vaughn, Rasmus Nielsen

**Author notes:** University of California, Berkeley Department of Integrative Biology 3040 Valley Life Sciences Building #3140 Berkeley, CA 94720-3140.

## Abstract

The ancestral recombination graph (ARG) is a structure that describes the joint genealogies of sampled DNA sequences along the genome. Recent computational methods have made impressive progress towards scalably estimating whole-genome genealogies. In addition to inferring the ARG, some of these methods can also provide ARGs sampled from a defined posterior distribution. Obtaining good samples of ARGs is crucial for quantifying statistical uncertainty and for estimating population genetic parameters such as effective population size, mutation rate, and allele age. Here, we use standard neutral coalescent simulations to benchmark the estimates of pairwise coalescence times from three popular ARG inference programs: ARGweaver, Relate, and tsinfer+tsdate. We compare 1) the true coalescence times to the inferred times at each locus; 2) the distribution of coalescence times across all loci to the expected exponential distribution; 3) whether the sampled coalescence times have the properties expected of a valid posterior distribution. We find that inferred coalescence times at each locus are most accurate in ARGweaver, and often more accurate in Relate than in tsinfer+tsdate. However, all three methods tend to overestimate small coalescence times and underestimate large ones. Lastly, the posterior distribution of ARGweaver is closer to the expected posterior distribution than Relate’s, but this higher accuracy comes at a substantial trade-off in scalability. The best choice of method will depend on the number and length of input sequences and on the goal of downstream analyses, and we provide guidelines for the best practices.

---

The full ARG is a structure that encodes all coalescence and recombination events resulting from the stochastic process of the coalescent with recombination. Hudson (1983) first described a stochastic process that combines recombination and coalescence to generate genealogies. At each given site, the genealogy resulting from this process is equivalent to the one generated by the single-locus coalescent model (Kingman 1982), but because recombination breaks loci apart (Figure 1A), the local genealogies can differ between sites.

**Figure 1.**
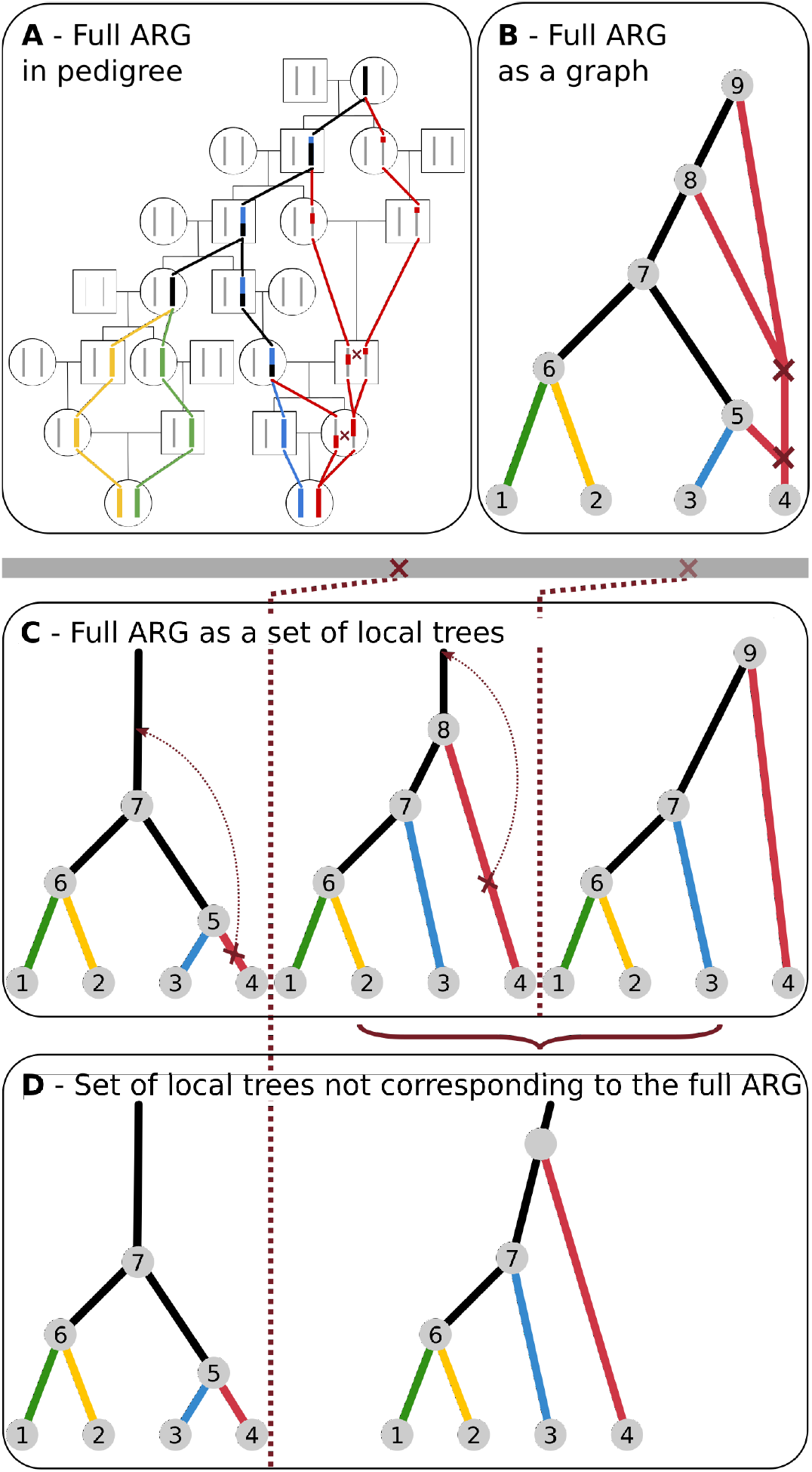
Schematic representations of the genealogy of a sample of two diploid individuals. Colors denote the four haplotypes sampled, and black lines indicate lineages or sequence tracts where at least one coalescence has occurred. Dark red crosses indicate recombination events. (A) The genealogy embedded in a pedigree. (B) An ancestral recombination graph (ARG) that fully represents all genealogical relationships shown in A, assuming that recombination events are annotated with the sequence coordinates. (C) An equivalent representation of the full ARG as a set of local trees separated by a single recombination event. (D) A set of trees that does not correspond to the full ARG. Instead, the second tree is an average of the local trees at that region.This set of trees is missing a recombination event that does not change topology, but changes the coalescence time. Other types of recombination events that could me missing in a partial ARG are: i) recombination followed by coalescence in the same branch, which does not change topology or other coalescence times and ii) topology changing recombination events.

## Representations of the ARG

The full ARG can be represented as a directed graph with two types of nodes: 1) coalescence nodes, where two or more edges merge into one (backwards in time) and 2) recombination nodes, where one edge splits in two (backwards in time) (Figure 1B). Alternatively, the full ARG can also be represented as an ordered collection of marginal coalescence trees, annotated with the recombination nodes. These marginal trees are embedded in the graph representation (Figure 1B,C).

In some representations, the collection of trees may or may not contain all the information from the full ARG, depending on whether the times of recombination events (red crosses in Figure 1) are stored with the trees (Rasmussen *et al*. 2014), and whether the internal nodes of the tree are labelled so they can be explicitly shared between adjacent trees. Furthermore, in some cases only topology changing recombination events are represented, and thus information regarding recombination events that do not lead to topology changes can be lost (Kelleher *et al*. 2019). Finally, some representations of ARGs as a collection of local trees allow more than one recombination event between trees (Speidel *et al*. 2019). In the latter two cases, each tree will potentially be an average of multiple coalescence trees. Figure 1D shows an example of a collection of local trees that does not correspond to the underlying full ARG, since one of its local trees is an average of two adjacent trees with identical topologies.

Collections of local trees with labelled internal nodes, regardless of whether they represent a full ARG or not, can be represented efficiently in computer memory by noting that each branch is part of many marginal trees (note repeated node numbers across trees in Figure 1C)). This property has been explored in the “tree sequence” format (Kelleher *et al*. 2018).

The full ARG contains all the information in a sample of DNA sequences regarding demography. Specifically, for a set of demographic parameters *θ*, parameters of the mutational process *µ*, sequence data *x*, and ARG *G, p*(*x*|*θ, µ, G*) = *p*(*x*|*µ, G*), *i*.*e*. if *G* is known there is no more information in the data about *θ*. A similar statement can be made for recombination and selection, if the leaf nodes of *G* are augmented with the allelic state at the selected loci. Therefore, the ARG is necessarily at least as informative as the combination of any and all summary statistics traditionally used to infer evolutionary processes (such as *F*_*ST*_, *π*, Tajima’s *D*, or EHH). Knowledge of the ARG is key for constructing powerful methods for extracting population genetic information from DNA sequencing data.

### Inferring ARGs

Unfortunately, ARGs cannot be directly observed but must be inferred from the data. Together with an estimate of the ARG, it is desirable to quantify the uncertainty around the inferred ARG, for example by obtaining samples of ARGs according to their posterior probabilities under a given model (we discuss examples of these models in the next section). Such samples can be used to quantify uncertainty regarding ARG inferences in downstream analyses. Accurate sampling from the posterior distribution is especially relevant for downstream methods that rely on importance sampling to infer evolutionary parameters from ARGs. In essence, these methods weight parameter inference under each sampled ARG by the ARG probability and therefore require that the samples of ARGs accurately reflect their probability distribution. These types of methods can be used to infer population size history, selection (Stern *et al*. 2019), migration (Osmond and Coop 2021), mutation rates and recombination rates.

Inferring full ARGs and quantifying inference uncertainty by sampling from the posterior distribution is a challenging problem computationally. It requires navigating a high-dimensional distribution of ARGs, which are themselves a complicated data structure. For this reason, inferring ARGs and sampling from their posterior distribution seemed like a nearly impossible endeavour some years ago, but important methodological developments now allow us to do so. Today, there are several methods available to estimate the full ARG or approximations of it, including ARGweaver (Rasmussen *et al*. 2014), Relate (Speidel *et al*. 2019) and tsinfer+tsdate (Kelleher *et al*. 2019; Wohns *et al*. 2022).

### Approximations of the coalescent with recombination

The classical way to include recombination in coalescence models is to consider the temporal process of lineage splitting caused by recombination and lineage merger caused by coalescences as one moves backwards in time (Hudson 1983; Griffiths and Marjoram 1997) (Figure 1A,B). Wiuf and Hein (1999) considered instead the spatial process of recombination along a sequence. In this formulation, the ARG is constructed as a sequence of local coalescent trees along a genome, where each tree is separated from adjacent trees by recombination events (Figure 1C). At each recombination breakpoint, a new tree is formed from the immediately preceding tree. To form the next tree, first one of the branches in the current tree is detached. Next, a point earlier than the detachment point is randomly chosen from any of the branches in **any of the previous trees** in the sequence. Finally, the detached branch coalesces to this chosen point.

To improve the computational efficiency in simulations, McVean and Cardin (2005) proposed approximating the spatial process as a Markovian process called the Sequentially Markovian Coalescent (SMC). In the SMC, when a lineage is detached from a tree at a recombination event, it can only coalesce back to one of the other lineages present **at the current tree**. Marjoram and Wall (2006) proposed an improved approximation, the SMC’, in which the detached lineage can coalesce to any branch in the current tree, including the one it was detached from. This means that some recombination events in this model do not generate a different local coalescent tree. This simple modification significantly improves the model in terms of approximating the full coalescent (Marjoram and Wall 2006; Wilton et al. 2015).

A heuristic approximation to the coalescence with recombination proposed by Li and Stephens (2003), extending ideas from Stephens and Donnelly (2000), approximates the coalescent with recombination using a copying process where one sequence is modeled as a copy of other sequences in the sample, with errors representing mutations and switches in the copying template representing recombination events. While this model has disadvantages, such as a dependence on the input order of sequences, it has proven computationally convenient for many purposes, including demography inference, introgression detection, and more (Sheehan et al. 2013; Steinrücken et al. 2018, 2019).

The formulation of the coalescent with recombination approximated as a Markovian process generating tree sequences in the SMC (McVean and Cardin 2005) and SMC’ (Marjoram and Wall 2006) and as a copying process of individual sequences by Li and Stephens (2003), paved the way for more scalable ARG inference methods. Notably, ARGweaver (Rasmussen et al. 2014) based on the SMC or SMC’ model, and Relate (Speidel et al. 2019) and tsinfer+tsdate (Kelleher et al. 2019; Wohns et al. 2022) based on the model by Li and Stephens (2003).

### ARGweaver

ARGweaver uses Markov Chain Monte Carlo (MCMC) to sample ARGs from the posterior distribution under the SMC or SMC’. It relies on a discretization of time (such that all recombination and coalescence events are only allowed to happen at a discrete set of time points) which makes the state space of ARGs finite countable and allows the use of discrete state-space Hidden Markov Models (HMMs). It then uses a lineage threading approach, which is a Gibbs sampling update, to sample the history of a single lineage or haplotype from the full conditional posterior distribution given the rest of the ARG connecting all other haplotypes.

### Relate

Relate simplifies the problem of ARG inference by inferring marginal coalescence trees, instead of full ARGs. Inference is divided into 2 steps. First, the Li and Stephens (2003) haplotype copying model is used to calculate pairwise distances between samples in order to infer local tree topologies. Next, it uses MCMC under a coalescent prior to infer coalescence times on those local trees. Relate is able to output samples of coalescence times from the posterior distribution using this MCMC approach, but it does so for the same fixed sequence of tree topologies. This is different from the ARGweaver MCMC sampling, which also samples the tree topology space (Table 1).

**Table 1.** Genome-wide genealogy inference programs compared.

| Program | Samples<br>topologies | Samples<br>coalescence times | Supports<br>demographic model | Scalability<br>(number of genomes) | Outputs<br>full ARG | Supports<br>unphased data |
| --- | --- | --- | --- | --- | --- | --- |
| ARGweaver | Yes | Yes | No | $\sim 50$ | Yes | Yes |
| ARGweaver-D* | Yes | Yes | Yes | $\sim 50$ | Yes | Yes |
| Relate | No | Yes | No | $\sim 10^3$ | No | No |
| tsinfer+tsdate | No | No | No | $\sim 10^5$ | No | No |

\* Hubisz *et al.* (2020)

### tsinfer, tsdate, and the tree sequence framework

Tsdate (Wohns *et al*. 2022) is a method that estimates coalescence times of tree sequences. Here, we used this method to date tree sequences inferred by tsinfer (Kelleher *et al*. 2019). Similarly to Relate, tsinfer is also based on the copying process from Li and Stephens (2003). A key innovation of tsinfer is a highly efficient tree sequence data structure which stores sequence data and genealogies (Kelleher *et al*. 2016, 2018, 2019; Ralph *et al*. 2020). Tsinfer performs inference in two steps. First, it recreates ancestral haplotypes based on allele sharing between samples. Next, it uses an HMM to infer the closest matches between ancestral haplotypes and the sampled haplotypes using an ancestral copying process modified from the classical Li and Stephens (2003) model to generate the tree topology. Finally, nodes in tree sequences inferred by tsinfer can be dated by tsdate. Tsdate uses a conditional coalescent prior, where the standard coalescent is conditioned on the number of descendants of each node on a local tree. Like ARGweaver, tsdate also discretizes time for computational efficiency. This framework infers a fixed topology and coalescence time, but it has the potential to sample coalescence times.

### Benchmarking of ARG inference methods

Here, we use standard neutral coalescent simulations to benchmark coalescence time inferences in ARGweaver (Rasmussen *et al*. 2014), Relate (Speidel *et al*. 2019), and tsinfer+tsdate (Kelleher *et al*. 2019; Wohns *et al*. 2022). We focus mainly on ARGweaver and Relate because they report measures of uncertainty in inference by allowing the user to output multiple samples from the posterior distribution. Sampling from the posterior is not currently implemented in tsdate (Table 1), but we include it in this evaluation because it is a promising framework for very fast tree-sequence inference, and it will likely provide an option to output samples from the posterior distribution of tree-sequences in future updates.

We focus our analyses on coalescence times not only because they are a very informative statistic about evolutionary processes, but also because they can be fairly compared across all methods. More specifically, ARGweaver and tsdate allow for polytomies (*i*.*e*., more than two branches coalesce at the same node). Relate, on the other hand, does not allow polytomies. Comparing topologies with and without polytomies could bias our results depending on how we chose to deal with polytomies, so we decided to focus on coalescence times only.

We run coalescent simulations on msprime (Kelleher *et al*. 2016) and compare the true (simulated) ARGs to the ARGs inferred by ARGweaver, Relate, and tsinfer+tsdate. We compare the ARGs with respect to their pairwise coalescence times using three different types of evaluation (Figure 2). First, we compare the true pairwise coalescence time at each site to the inferred time. Second, we compare the overall distribution of pairwise coalescence times across all sites and all MCMC samples to the expected distribution. In Bayesian inference, the data averaged posterior distribution is equal to the prior. Since data are simulated under the standard coalescent with recombination the data averaged posterior should be exponential with rate 1 in coalescence time units (2*N*_*e*_ generations, where *N*_*e*_ is the effective population size). Third, we used simulation-based calibration (SBC) (Cook et al. 2006; Talts et al. 2020) to evaluate if the posterior distributions sampled by ARGweaver and Relate are well calibrated (see details in Methods).

**Figure 2.**
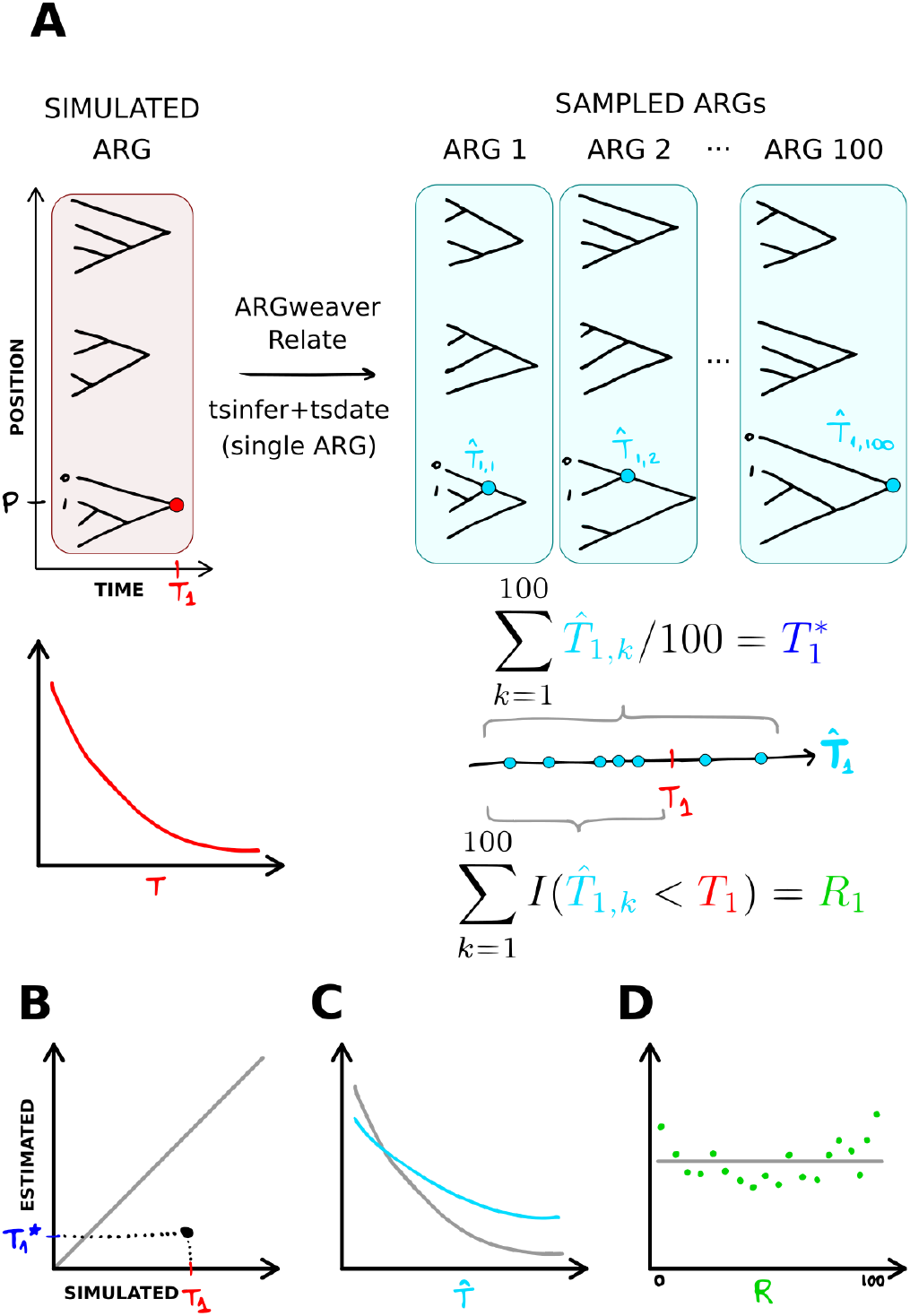
Methods overview. (A) Data (ARGs and DNA sequences) were simulated from the coalescent with recombination. In the model and simulated data, pairwise coalescence times (CT) are exponentially distributed (Figure S3). *T*_1_ represents the CT between samples 0 and 1, at position P in the simulated data. 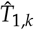 is the CT between samples 0 and 1 at position P, in each ARG sample *k*. Point estimates 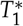 are obtained as the mean of 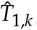, and the rank statistic is computed as the number of 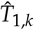 that are smaller than the true value *T*_1_. (B) We compare estimated to simulated values of the CT of each pair of samples, at each position of the genome. (C) We compare the distribution of sampled CT across all sampled ARGs, all sites and all pairs of samples to the expected exponential distribution. (D) We compare the distribution of ranks to the expected uniform distribution.

## Methods

### Simulations

We simulated tree sequences and SNP data with msprime version 0.7.4 (Kelleher et al. 2016). For simulations with Jukes and Cantor (1969) mutational model, we used msprime version 1.0.2 (Baumdicker et al. 2021) to add mutations to trees simulated under msprime 0.7.4, because the Jukes and Cantor (1969) model option was not available in msprime 0.7.4. Unless otherwise noted, simulations were done under the standard neutral coalescent (Hudson model in msprime) and using the following parameters: 4 diploid samples (*i*.*e*. 8 haplotypes), total map length *R* = 20000 and mutation to recombination rate ratio *µ*/*ρ* = 1. In practice, we used the following parameter values in msprime: effective population size of 10,000 diploids (2*N*_*e*_ = 20, 000), mutation rate and recombination rate of 2 × 10^−8^ per base pair per generation and a total sequence length of 100Mb.

We varied these standard simulation scenarios in several ways: using SMC and SMC’ models, different numbers of samples (4, 16, 32 and 80 haplotypes), a 10-fold increase and 10-fold decrease in the mutation to recombination ratio (in each case changing either the mutation or the recombination rates), and changing the total length of input sequence from 100Mb to 5Mb and 250kb. These simulated sequences were then divided into 20 equally sized segments, so that ARGweaver could be run on each in parallel (see below). The minimum length of total simulated sequence (250kb) was chosen such that the average number of pairwise differences between each of the 20 segments was 10, given a mutation rate of 2 *×* 10^−8^.

We extracted coalescence times at all sites in the simulated trees in BED format (columns: chromosome, start position, end position, coalescence time), with one BED file for each pair of samples. Figure 2 shows an overview of the metrics extracted from simulated ARGs and from ARGs estimated by tsinfer+tsdate or sampled from the posterior by ARGweaver and Relate.

### ARGweaver

VCF files from msprime were converted to ARGweaver sites format using a custom python script. We ran ARGweaver’s *arg-sample* program to sample ARGs. This was done in parallel on 20 segments of equal size, using the *–region* option. We used the same values used in the msprime simulations (*–mutrate* and *–recombrate 2e-8* and *–popsize 10000*) and except where otherwise noted, we ran ARGweaver using the SMC’ model (*–smcprime* option). We ran ARGweaver with 1200 or 2200 iterations (*–iters*) (with burn-in of the first 200 or 1200 iterations, respectively), depending on how long it took to converge. Assessment of convergence is described below and in the Supplementary Materials, Evaluating MCMC Convergence. We extracted 100 MCMC samples from every 10th iteration among the last 1,000 iterations (default *–sample-step 10*).

We extracted all pairwise coalescence times in BED format with the program *arg-summarize* using options *–tmrca* and *–subset*, and we used bedops (version 2.4.35 (Neph et al. 2012)) to match the times sampled by ARGweaver to the simulated ones at each sequence segment. Finally, we used a custom Python script to calculate the ranks of simulated pairwise coalescence times on ARGweaver MCMC samples per site.

### Time discretization

In ARGweaver, time is discretized such that recombination and coalescence events are only allowed to happen at a user-defined number of time points, *K* (default value is 20) (Rasmussen *et al*. 2014). These time points *s*_*j*_ (for 0 *<*= *j <*= *K* − 1) are given by the function

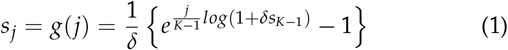

where *δ* is a parameter determining the degree of clustering of points in recent times. Small values of *δ* lead to a distribution of points that is closer to uniform between 0 and *s*_*K* −1_, and higher values increase the density of points at recent times (default value is 0.01) (Hubisz and Siepel 2020). Equation 1 ensures that *s*_0_ is always 0, and *s*_*K*−1_ (or *s*_*max*_) is user defined by the parameter *–maxtime* (default value is 200,000).

Rounding of continuous times into these *K* time points is done by defining bins with breakpoints between them, such that the breakpoint between times *s*_*j*_ and *s*_*j*+1_ is 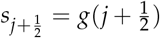. All continuous values in the bin between 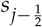 and 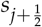 are assigned the value *s*_*j*_. We note that for the first and last intervals, the values assigned (*s*_0_ and *s*_*K*−1_) do not correspond to a midpoint in the time interval but rather to its minimum (*s*_0_ = 0) or maximum (*s*_*K*−1_ = *s*_*max*_)

Here, when reporting results in bins, we use the same time discretization as defined by the ARGweaver breakpoints 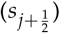. However, we change the value assigned to times in these bins: instead of using *s*_*j*_, we define *t*_*j*_ as the median of the exponential distribution with rate 1 at the interval between 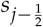 and 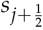. To this end, we first calculate the cumulative probability of the exponential distribution with rate 1 up to the median of the *j*th interval

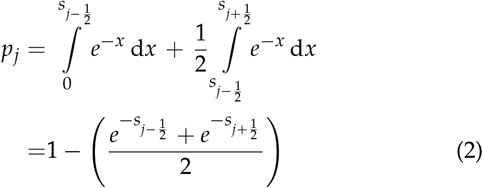

We then take the inverse CDF of the exponential distribution with rate 1, at the point *p*_*j*_, to find the time *t*_*j*_ = −*ln*(1 − *p*_*j*_) corresponding to the median value for the interval.

This step is relevant for the simulation-based calibration (see below), where we take the rank of true (simulated) coalescence times relative to the values sampled by ARGweaver. If we used *s*_*j*_, coalescence times in the first or last ARGweaver time interval would not be represented by a midpoint. We correct for that by using *t*_*j*_, so that all time intervals are comparable.

Relate does not use time discretization, and tsdate uses a discretization scheme where the time points are the quantiles of the lognormal prior distribution on node ages (Wohns *et al*. 2022). Here, we always apply the ARGweaver time discretization scheme when comparing results in bins.

### Relate

VCF files generated with msprime were converted to Relate haps and sample files using *RelateFileFormats –mode ConvertFromVcf* and Relate’s *PrepareInputFiles* script. We ran Relate (version 1.1.2) using *–mode All* with the same mutation rate (*-m 2e-8*) and effective population size (*-N 20000*) used in the msprime simulations, as well as a recombination map with constant recombination rate along the genome, with the same rate used in msprime (*2e-8*).

We used Relate’s *SampleBranchLengths* program to obtain 1000 MCMC samples of coalescence times for the local trees inferred in the previous step in anc/mut output format (*–num-samples 1000 –format a*). Similarly to the ARGweaver analysis, we also performed this step in 20 sequence segments of 5Mb, and we thinned the results to keep only every 10th MCMC sample. Finally, we extract pairwise coalescence times and calculate the ranks of true pairwise coalescence times relative to the 100 MCMC samples. Due to the large number of pairwise coalescence times, for the simulations with 80 and 200 samples, we extracted coalescence times from a subset of 210 pairs of samples. We extracted coalescence times for every 4th vs. every 4+1th sample in the case of 80 samples, and 10th vs. every 10+1th sample in the case of n=200.

### tsinfer and tsdate

VCF files generated by msprime were provided as input to the python API using *cyvcf2*.*VCF* and converted to tsinfer *samples* input object using the *add_diploid_sites* function described in the tsinfer tutorial (https://tsinfer.readthedocs.io/en/latest/tutorial.html#reading-a-vcf). Genealogies were inferred with tsinfer (version 0.2.0 (Kelleher *et al*. 2019)) with default settings and dated with tsdate (version 0.1.3 (Wohns *et al*. 2022)) using the same parameter values as in the simulations (*Ne=10000, mutation_rate=2e-8*), with a prior grid of 20 timepoints.

Pairwise coalescence times were extracted from the tree sequences using the function *tmrca()* from tskit (version 0.3.4 (Kelleher *et al*. 2018)), and output in BED format, with one file for each pair of samples. Finally, coalescence times at each site, for each pair of samples were matched to the simulated ones (also in BED format) using bedops (Neph *et al*. 2012).

### MCMC convergence

We evaluated MCMC convergence of Relate and ARGweaver through 1) visual inspection of trace plots, 2) autocorrelation plots, 3) effective sample sizes and 4) the Gelman-Rubin convergence diagnostics based on potential scale reduction factor (Gelman and Rubin 1992; Brooks and Gelman 1998). Trace plots were also used to determine the number of burn-in samples, and autocorrelation plots were used to determine thinning of the samples. See Evaluating MCMC Convergence in Supplementary Materials for details.

### Point estimates of pairwise coalescence times

We estimated pairwise coalescence times from the MCMC samples from Relate and ARGweaver by taking the average of 100 samples at each site (Figure 2). Since tsdate does not output multiple samples of node times, we use its point estimate of pairwise coalescence times directly. Point estimates of coalescence times were compared to the simulated values for each pair of samples, at each site along the sequence.

Mean squared error (MSE) of point estimates was calculated from each point estimate of coalescence time (for each pair of samples, at each site), as well as per bin of size 0.1 of the simulated coalescence times (in units of 2*N*_*e*_ generations) for Figure S2. We also report Spearman’s rank correlation (*r*_2_) of the point estimates of pairwise times in each tree against the simulated tree, averaged over all positions in the genome.

### Simulation-based calibration

In addition to comparing MCMC point estimates to the true simulated values, we use simulation methods proposed by Cook *et al*. (2006) and Talts *et al*. (2020) to assess whether Bayesian methods are sampling correctly from the true posterior distribution. Cook *et al*. (2006) proposed simulating data using parameters sampled from the prior. The posterior, when averaged over multiple simulated data sets, should then equal the prior.

In our case, we sample ARGs, *G*, from the full coalescence process with recombination with a known implicit prior of pairwise coalescence times, *P*(*t*) = *e*^−*t*^. We simultaneously simulate sequence data, *x*, on the simulated ARGs from the distribution *p*(*x*) =∫ *p*(*x*|*G*)d*P*(*G*). The distribution of the averaged posterior of *G, p*_*ave*_ (*G*) = ∫ *p*(*G*|*X*)d*P*(*x*) should then equal the prior for *G* (Talts et al. 2020), and hence the prior distribution for the pairwise coalescence times, *t*, should equal the averaged posterior distribution for *t*. Here, all population parameters relating to mutation, effective population sizes, etc., are kept fixed and suppressed in the notation. One way we will examine the accuracy of the posterior inferences is, therefore, to compare the average of the posterior of *t* to the exponential distribution. In practice, we simulate data using msprime (Kelleher et al. 2016) and pipe the data to the MCMC samplers (ARGweaver and Relate) for inference of the posterior distribution. ARGweaver uses an approximation (SMC’) of the model (coalescent with recombination) used in the data simulations, and Relate uses a heuristic method based on the Li and Stephens model. Thus, inadequacies of the fit of the posteriors could potentially be caused by this discrepancy between the model used in simulations and the models used for inference.

However, even if the averaged posterior resembles an exponential, the inferences for any particular value of *t* may have a posterior that is too narrow or too broad. For a closer examination of the accuracy of the posterior, we use a method proposed by Cook *et al*. (2006) and Talts *et al*. (2020) that compares each posterior to the true value. To this end, we compare each true (simulated) pairwise coalescence time to the corresponding posterior for the same pair of haplotypes. If the posterior is correctly calculated, the rank of the true value relative to the samples from the posterior should be uniformly distributed (Cook *et al*. (2006), Talts *et al*. (2020)). We use 100 MCMC samples from ARG-weaver and Relate for each data set, meaning our ranks take values from 0 to 100. Deviations from the uniform distribution of ranks quantifies inaccuracies in estimation of the posterior. For example, an excess of low and high ranks indicates that the inferred posterior distribution is underdispersed relative to the true posterior.

## Results

### Comparison of simulated to estimated coalescence time per site

We compared coalescence times estimated by ARGweaver, Relate and tsinfer+tsdate to the true values known from msprime simulations. In all three methods, estimates of coalescence time per site are biased (Figure 3 and S2). Small values of coalescence times are generally overestimated, while large values tend to be underestimated (Fig S2). In tsinfer+tsdate, point estimates are apparently bounded to a narrow range (Figure 3G). The mean squared error (MSE) of point estimates is larger in Relate (MSE=0.625) and tsinfer+tsdate (MSE=1.631) than in ARG-weaver (MSE=0.397), showing that point estimates of pairwise coalescence times at each site are closer to the true value in ARG-weaver. Spearman’s rank correlation is also highest in ARG-weaver (*r*_*s*_=0.761), but in this metric tsinfer+tsdate (*r*_*s*_=0.705) perform better than Relate (*r*_*s*_=0.669).

**Figure 3.**
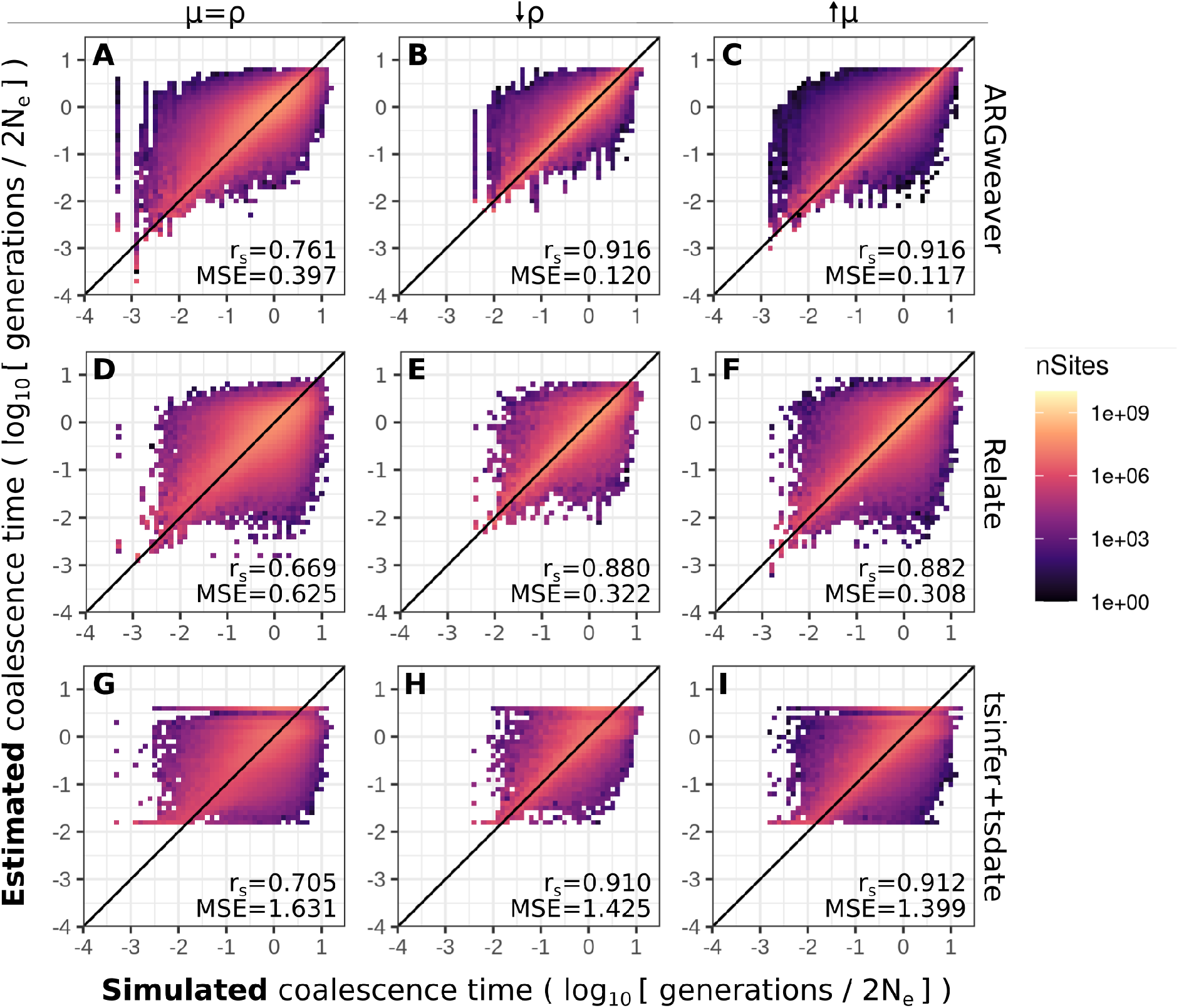
Point estimates of coalescence times in ARGweaver (A-C), Relate (D-F) and tsinfer+tsdate (G-I). Left column: *µ* = *ρ* = 2 *×* 10^−8^; middle column: *µ*/*ρ* = 10, *ρ* = 2 *×* 10^−9^; right column: *µ*/*ρ* = 10, *µ* = 2 *×* 10^−7^. For ARGweaver and Relate, point estimates are the means of 100 MCMC iterations. Note that axes are in log scale. See Figure S1 for the data in plots A,D,G plotted in linear axes. Diagonal line shows x=y. MSE: Mean squared error; *r*_*s*_: Spearman’s rank correlation.

For ARGweaver and Relate, the point estimates of coalescence times are obtained as the means of samples from the posterior. These Bayesian estimates are not designed to be unbiased and unbiasedness of the point estimator is arguably not an appropriate measure of performance for a Bayesian estimator. Therefore, we also evaluate the degree to which the posterior distributions reported by ARGweaver and Relate are well calibrated, *i*.*e*. represent distributions that can be interpreted as valid posteriors, and the degree to which the data-averaged posterior distributions of coalescence times equals the prior exponential distribution.

### Posterior distribution of coalescence times

We simulated data under the standard coalescent model, where the distribution of pairwise coalescence times (in units of 2*N*_*e*_ generations, where *N*_*e*_ is the diploid effective population size) follows an exponential distribution with rate parameter 1 (Figure S3). As argued in the Methods section, the same is true for the data-averaged posterior.

We compared the expected exponential distribution of coalescence times to the observed distribution of coalescence times across all sites inferred by ARGweaver, Relate, and tsinfer+tsdate (Figure 4). For ARGweaver and Relate, we output 100 MCMC samples from the posterior distribution and plot the distribution of pairwise coalescence times across all sites and MCMC samples.

**Figure 4.**
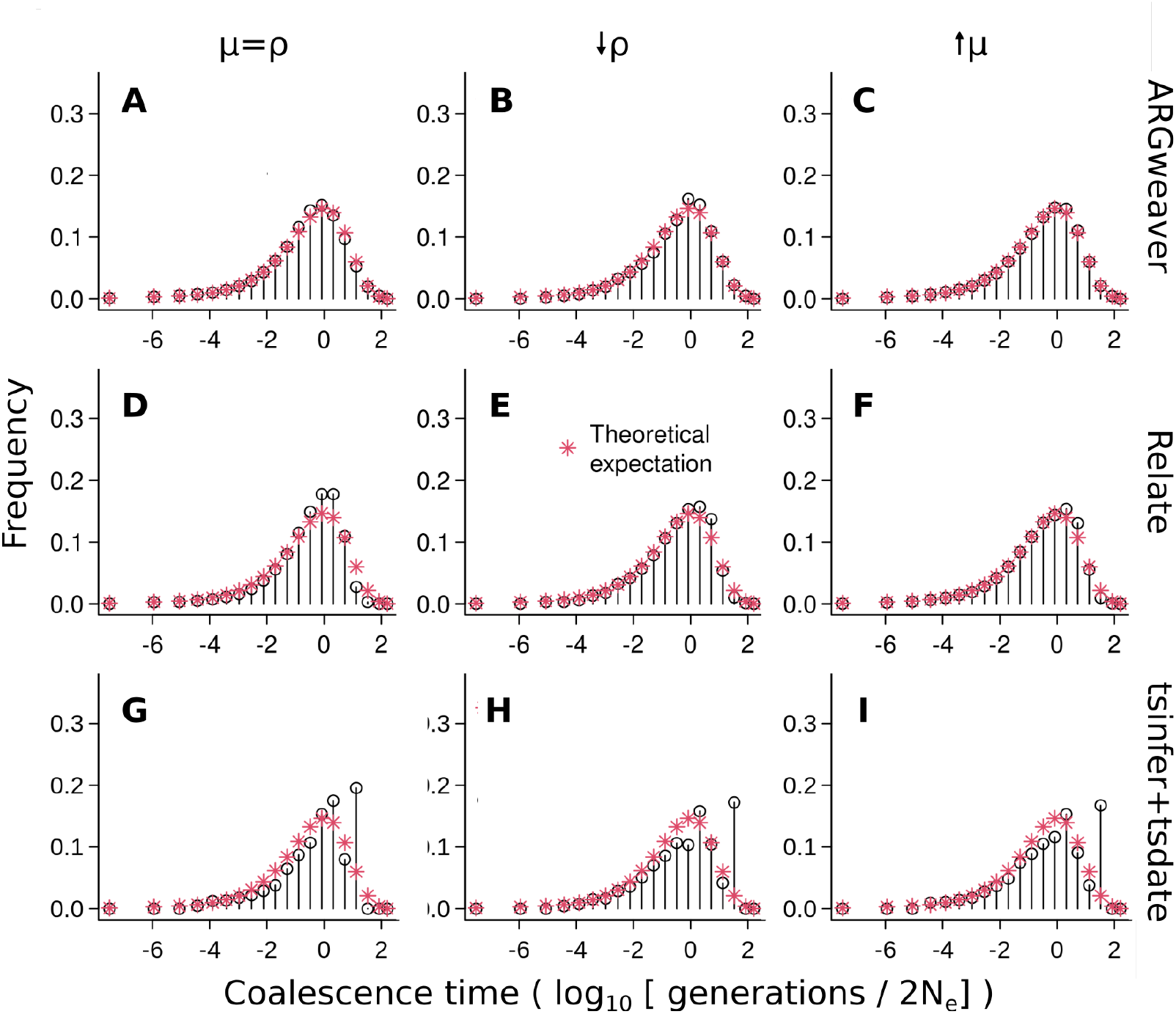
Distribution of coalescence times inferred by ARGweaver (A-C), Relate (D-F) and tsinfer+tsdate (G-I). Left column: *µ* = *ρ* = 2 ∗ 10^−8^; middle column: *µ*/*ρ* = 10, *ρ* = 2 ∗ 10^−9^; right column: *µ*/*ρ* = 10, *µ* = 2 ∗ 10^−7^. Plots D and G show the same data as in Figure S4, using different binning.

To facilitate visual comparison of the distributions between methods, we discretized Relate and tsinfer+tsdate coalescence times into the same bins as ARGweaver (Figure 4D,G, see distributions without discretization in Figure S4 and see Methods for a description of ARGweaver time discretization). Because the time discretization breakpoints are regularly spaced on a log scale, we use a log scale on the x-axis for better visualization.

Distributions of coalescence times from ARGweaver and Relate (Figure 4A and D) show an excess around 1, when compared to the expected exponential distribution. However, that bias is more pronounced in Relate than ARGweaver. In tsinfer+tsdate, the distribution is truncated at 1.6, and it deviates more strongly from the expected exponential distribution (Figure 4G). We note that the plots from ARGweaver and Relate are not directly comparable to those of tsinfer+tsdate, since there are 100 coalescence time samples at each site from the former two programs but only one from tsdate.

### Simulation-based calibration

In this section, we use simulation-based calibration to evaluate whether ARGweaver and Relate are generating samples from a valid posterior distribution of coalescence times (see Methods). To that end, we simulated coalescence times at multiple sites following the standard coalescent prior distribution, and we generated 100 MCMC samples from the posterior distribution using both ARGweaver and Relate. Finally, we analyse the distribution of the ranks of the simulated coalescence times relative to the 100 sampled values at each site.

In the previous section, we showed that the posterior distributions of ARGweaver and Relate are similar to the theoretically expected exponential distribution. However, in that analysis we have not evaluated the distribution of MCMC samples relative to each simulated value. The results of simulation-based calibration are informative about that distribution and can reveal if the posterior distribution is well calibrated.

The distribution of ranks from ARGweaver (Figure 5A, Kullback-Leibler Divergence (KLD) =0.027) is closer to uniform than that of Relate (Figure 5D, KLD=0.602). However, both show an excess of low and high ranks. The excess of low and high ranks indicates that the sampled posterior distribution is underdispersed (Talts et al. 2020), *i*.*e*. the posterior has too little variance and does not represent enough uncertainty regarding the coalescence times.

**Figure 5.**
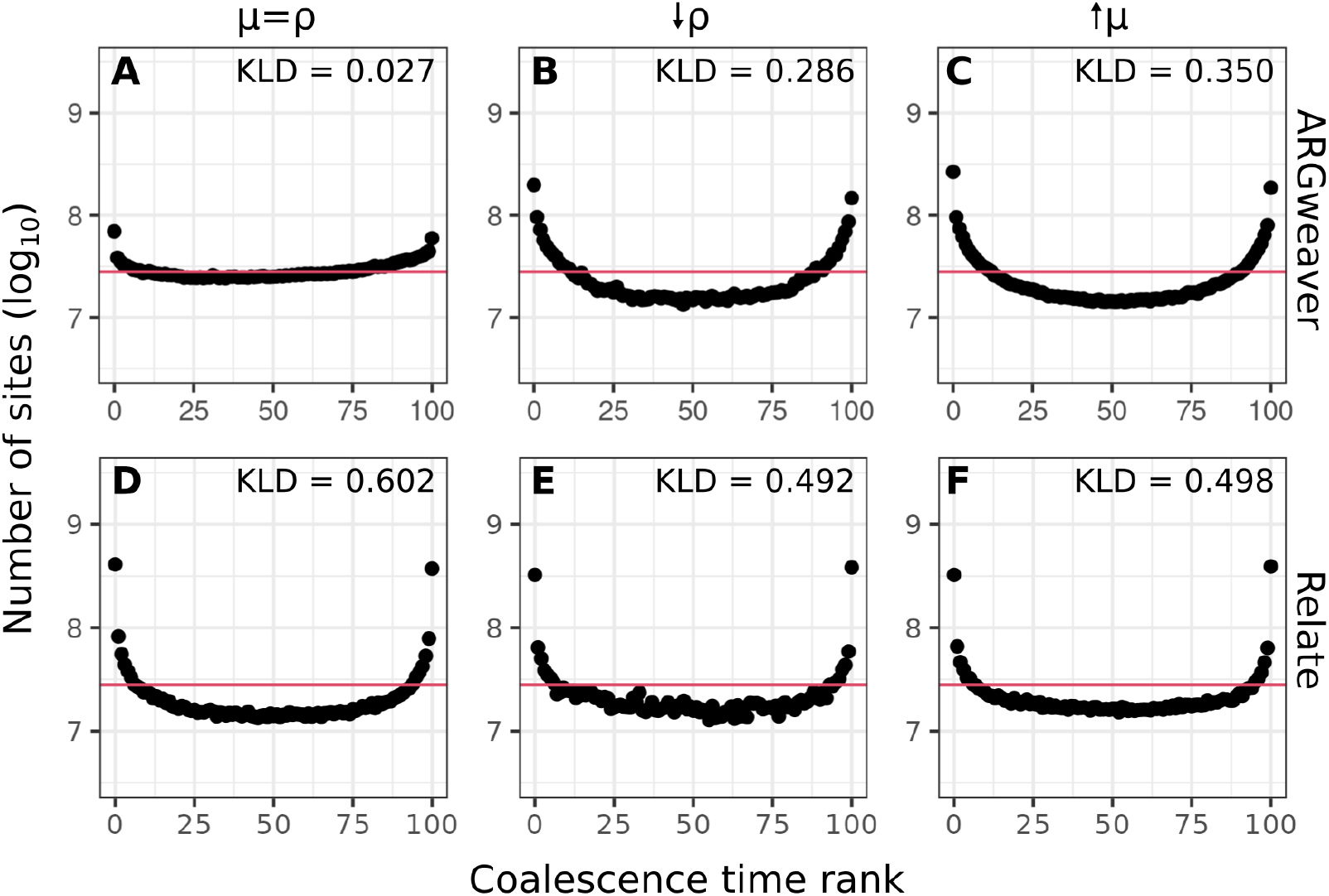
Counts of ranks from simulation-based calibration in ARGweaver (a-c) and Relate (d-f). Horizontal line shows expected uniform distribution. Left column: *µ* = *ρ* = 2 ∗ 10^−8^; middle column: *µ*/*ρ* = 10 decreasing recombination rate (*ρ* = 2 ∗ 10^−9^); right column: *µ*/*ρ* = 10 increasing mutation rate (*µ* = 2 ∗ 10^−7^). Horizontal line shows expected uniform distribution.

One possible cause for this type of deviation from the uniform distribution could be MCMC convergence, *i*.*e*., samples being autocorrelated, resulting in effective sample size is lower than the number of samples taken, the MCMC chain not mixing well and/or the MCMC chain not being run long enough to achieve convergence.

We show detailed results for MCMC convergence in Relate and ARGweaver in the Supplementary Materials. Briefly, we have not found these types of convergence issues in ARGweaver or Relate with simulations of 8 haplotypes and mutation to recombination ratio of 1. Potential scale reduction factor (PSRF) from Gelman-Rubin convergence diagnostic statistics are all close to 1 (Tables S2, S3), and effective sample sizes are almost all larger than 100. Therefore, MCMC convergence does not seem to explain why the rank distributions are not uniform.

### Increased mutation to recombination ratio

When inferring an ARG from sequence data, the information for inference comes from mutations that cause variable sites in the sequence data. The lower the recombination rate, the longer the span of local trees will be and the more mutations will be available to provide information about each local tree. More generally, an increased mutation to recombination ratio is expected to increase the amount of information available to infer the ARG.

In our standard simulations presented so far, the mutation to recombination ratio is one (*µ* = *ρ* = 2 ∗ 10^−8^). We increased the simulated mutation to recombination ratio to 10, both by decreasing the recombination rate (*ρ*) tenfold and also by increasing the mutation rate (*µ*) tenfold. We expected that these scenarios would improve inference of ARGs, and consequently the estimates of pairwise coalescence times. Point estimates are better with increased mutation to recombination ratio in ARG-weaver (Figure 3B,C), Relate (Figure 3E,F) and tsinfer+tsdate (Figure 3H,I).

The coalescence times distribution in Relate (Figure 4E,F) are closer to the expected with *µ*/*ρ* = 10 relative to *µ*/*ρ* = 1 (Fig 4D), and the simulation-based calibration also improved (Figure 5D-F, KLD=0.492 and 0.498 compared to KLD=0.602).

The results from ARGweaver with *µ*/*ρ* = 10 were more surprising, with the simulation-based calibration showing a more pronounced underdispersion of the posterior distribution (Figure 5B,C, KLD=0.286 and 0.350, compared with KLD=0.027 for *µ*/*ρ* = 1). The overall distribution of coalescence times, however, showed little change (Figure 4B,C). One possible explanation for ARGweaver results being worse with higher mutation to recombination ratio might be that MCMC mixing is worse under those conditions, leading to convergence issues not observed for the previous scenario. Examining convergence diagnostics seems to confirm this with more coalescence times showing low effective sample size, and with a potential scale reduction factor showing evidence of lack of convergence of some coalescence times (see Evaluating MCMC Convergence in Supplementary Materials).

We show additional simulation results in the Supplementary Materials, including simulations with reduced *µ*/*ρ*, which could be a realistic scenario around recombination hotspots (Figures S5 and S6) and ARGweaver results on simulations with intermediate values of *µ*/*ρ* (2 and 4), under the SMC and SMC’ genealogy models, and with the Jukes-Cantor mutation model in the Supplementary Materials.

### Number of samples

Next, we evaluate ARG inference with simulations with different sample sizes. Our standard sample size used so far was 8 haplotypes, and here we change it to 4, 16 and 32. For Relate and tsinfer+tsdate, which are scalable to larger sample sizes, we also evaluated inference with 80 and 200 sampled haplotypes.

For ARGweaver, increasing sample sizes decreased the MSE of point estimates (Figure 6A-C), distributions of coalescence times remained similar (Figure 7A-C), but underdispersion of the posterior distribution increased (Figure 8A-C). As mentioned in the previous section, this could be caused by an MCMC mixing problem. In particular, a larger number of samples will contribute to an increasing number of states for ARGweaver to explore, possibly leading to poor MCMC convergence (see Evaluating MCMC Convergence).

**Figure 6.**
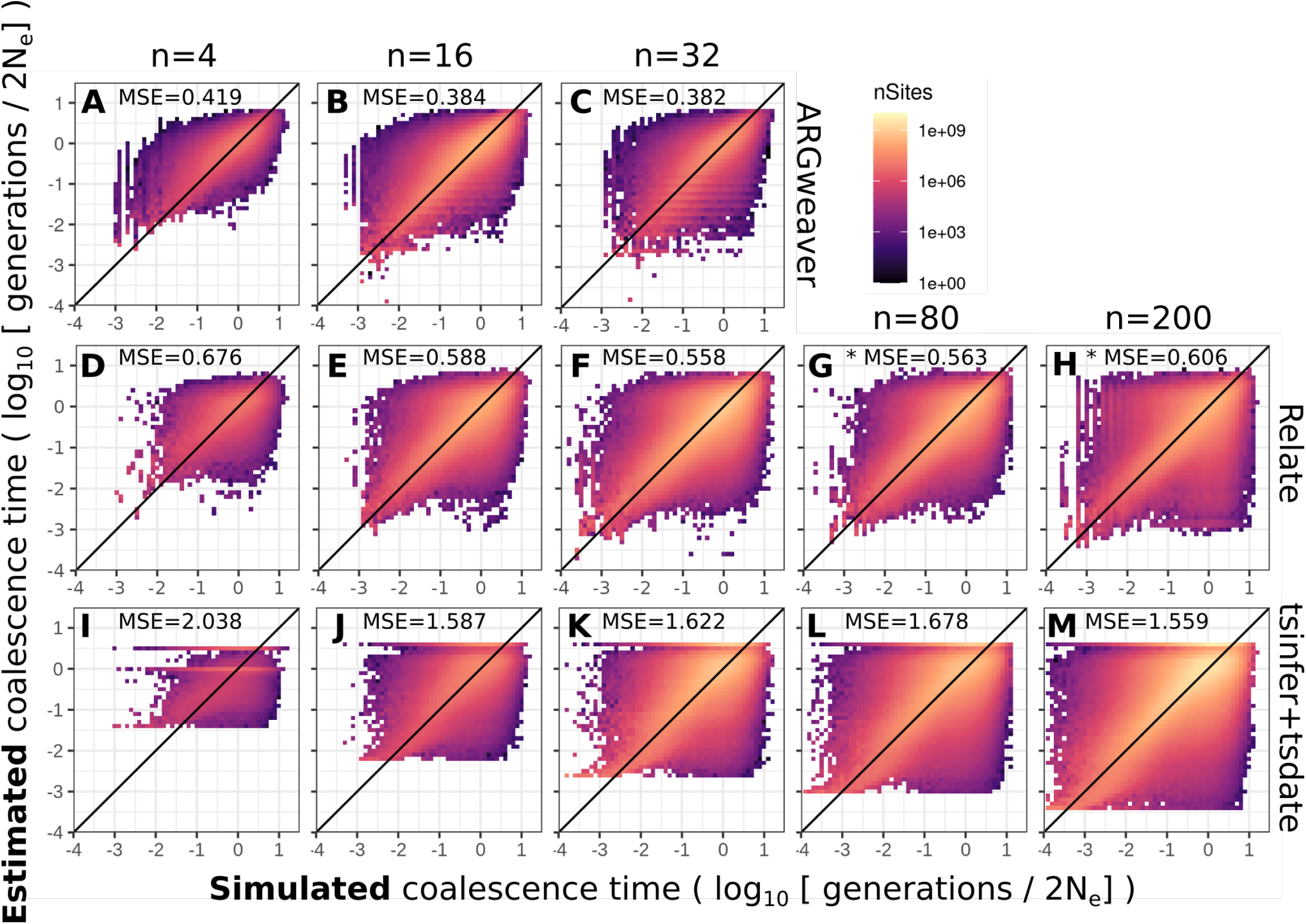
Point estimates of ARGweaver (A-C), Relate (D-H) and tsinfer+tsdate (I-M). Columns show different number of simulated samples 4, 16, 32, 80 or 200 haplotypes. Mean squared error (MSE) is shown for each plot. Note that ARGweaver is not scalable for simulations with larger sample sizes. * indicate results for a subset of 210 pairs of samples, instead of all pairwise coalescence times.

**Figure 7.**
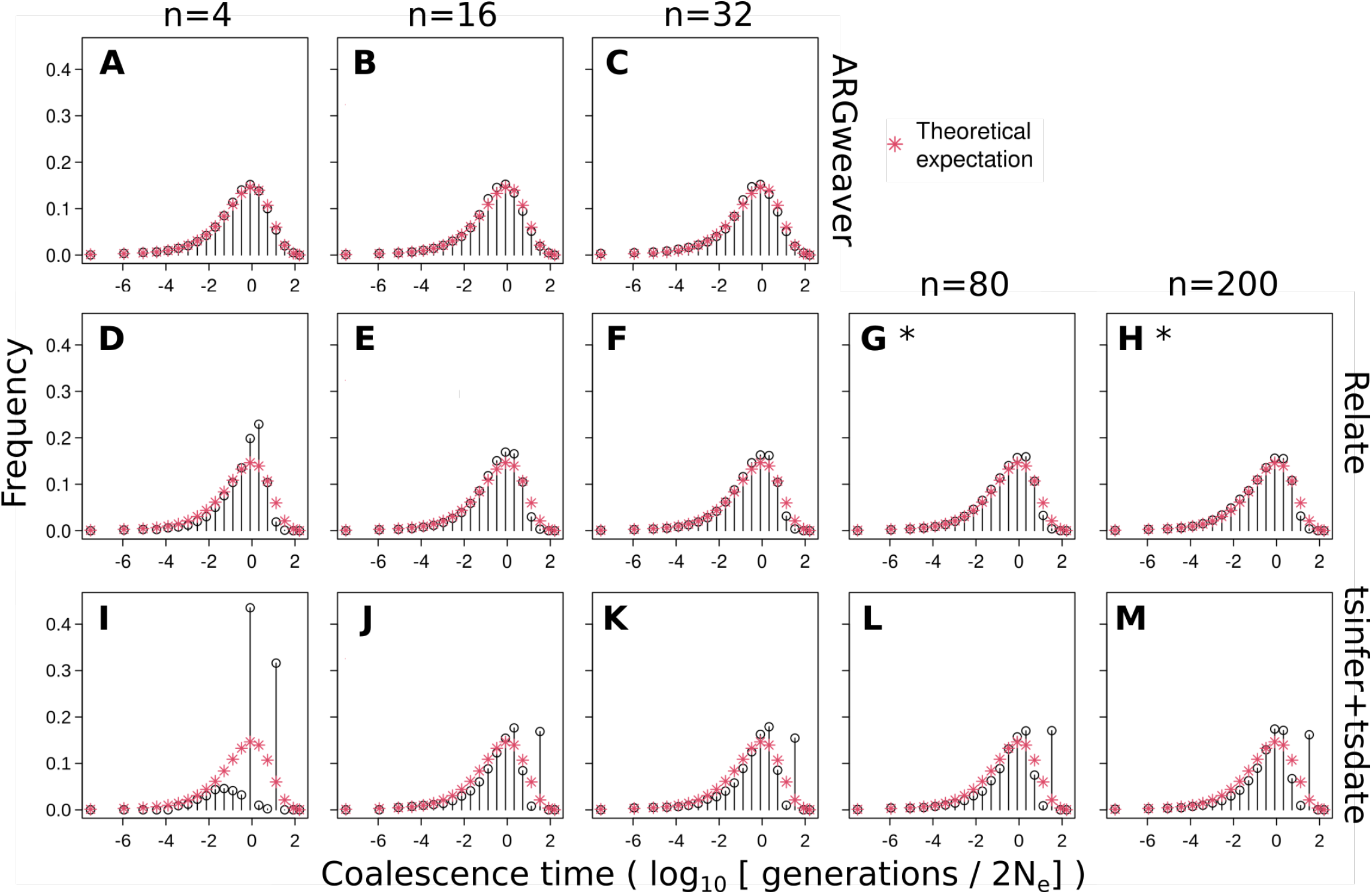
Distribution of coalescence times in ARGweaver (A-C), Relate (D-H) and tsinfer+tsdate (I-M). Columns: sample sizes of 4, 16, 32, 80, 200 haplotypes. * indicate results for a subset of 210 pairs of samples, instead of all pairwise coalescence times.

**Figure 8.**
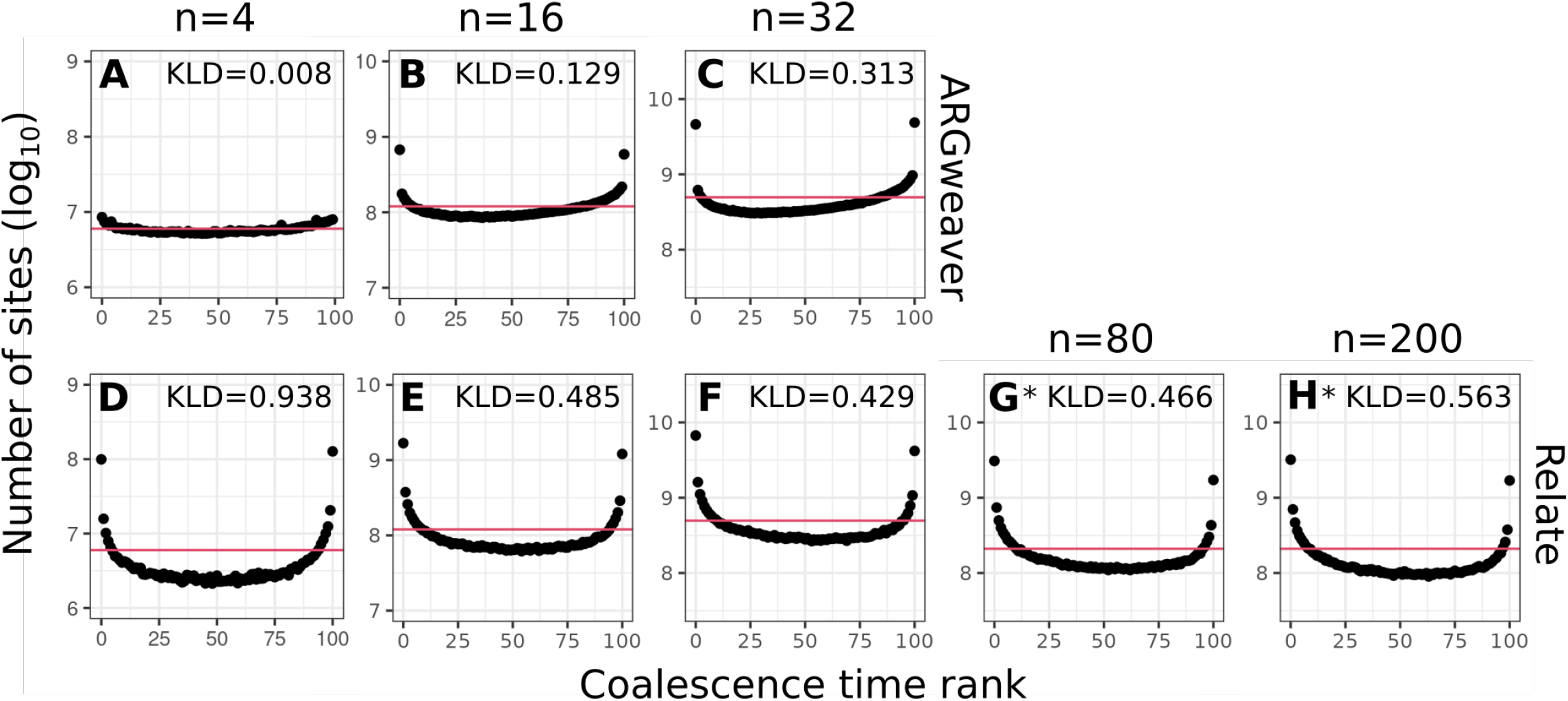
Simulation-based calibration for ARGweaver (A-C) and Relate (D-H). Columns: sample sizes of 4, 16, 32, 80, 200 haplotypes. Horizontal line shows expected uniform distribution. Note that the y-axis is centralized on different values but always has the same length. * indicate results for a subset of 210 pairs of samples, instead of all pairwise coalescence times.

With a smaller sample size (n=4 haplotypes), the coalescence time distribution from Relate showed an excess around the mean value (coalescence time of 1) (Fig 7D). With increasing sample sizes, it became more similar to the expected distribution (Fig 7E-H). Calibration of the posterior distribution improved with increasing sample sizes up to 32 haplotypes (Figure 8D-H).

Both the point estimates and posterior distribution of coalescence times in tsinfer+tsdate do not consistently improve or worsen with increasing sample sizes in the range tested here (Figures 6I-M and 7I-M).

### Length of input sequence

Point estimates of all programs remained similarly accurate when a much shorter input sequence was provided (5mb and 250kb, Figure S7A-C and S8A-C, compared with 100Mb in previous analyses). The distribution of coalescence times with 5Mb input sequence remained similar to the ones inferred with 100Mb input sequence (Figure S7D-F). However, distributions from simulations with only 250kb input sequence are visibly more deviated from the expected exponential distribution (Figure S8D-F). Distributions of ranks are noisier with decreasing input sequence length, but KLD remained similar (Figure S8 and S7H,G).

### Runtime

We point out that runtimes differ widely among the programs compared here, and this factor should be taken into account for users making decisions on what method to use for their applications. For example, in the simulations with mutation rate equal to recombination rate, with sample size of 8 haplotypes and taking 1000 MCMC samples, ARGweaver took a total of 641 computing hours while Relate took 17 hours. The clock time was reduced by running both programs in parallel for segments of 5Mb of the total 100Mb sequence, meaning that ARGweaver took approximately 35h. However, this still could be a significant amount of time for the user, depending on their utilization of the algorithm. For a systematic comparison of runtimes between Relate and ARGweaver, see Speidel *et al*. (2019). Impressively, tsinfer and tsdate took only 5 minutes.

## Discussion

ARG inference promises to be a tremendously useful tool for inferences of evolutionary history, such as natural selection or demography. However, it is also a very hard computational problem. We compared methods that use different approaches to this problem and evaluated their accuracy using simulated data and comparisons of three aspects of coalescence time estimates: 1) individual point estimates of each pairwise coalescence time; 2) the overall distribution of coalescence times across all sites; 3) the calibration of the reported posterior distributions.

Ancestral recombination graphs are extremely rich in information, including topological information of individual coalescence trees and information regarding the distribution of recombination events. We have not evaluated these aspects of inferred ARGs but have instead only focused on pairwise coalescence times. However, pairwise coalescence times are extremely informative statistics about many population-level processes and pairwise relationships between individuals, and they are also indirectly informative about tree topologies. Other research has compared the accuracy of tree topology inference (Rasmussen et al. 2014; Kelleher et al. 2019) and recombination rates (Deng et al. 2021) among ARG inference methods. We opted to focus on coalescence times not only because they are a very informative statistic about evolutionary processes, but also because they can be fairly compared across all methods. As described in the Introduction, comparisons of tree topologies could be confounded by the presence of polytomies in ARGweaver and tsinfer+tsdate and the absence of polytomies in Relate.

We found a strong speed-accuracy trade-off in ARG inference. ARGweaver performs best in our three tests: point estimates, the overall distribution of coalescence times, and the quality of sampling from the posterior. Importantly, it is also the only method we compared that resamples both topologies and node times (Table 1). This likely leads to a better exploration of ARG space and is one reason why it provides better samples from the posterior. On the other hand, it also contributes to making ARG-weaver much slower than the other methods and not scalable for genome-wide inference of 50 or more genomes.

Relate largely undersamples tree topologies (Deng *et al*. 2021), and thus every marginal tree estimate is only as good as an average over a series of true trees (Figure 1D). This will naturally lead to a more centered, under-dispersed distribution, as shown by the larger deviations from the uniform distribution in simulation-based calibration (Figures 5 and 8, where ARGweaver KLD values range from 0.008 to 0.350, and Relate range from 0.429 to 0.938). Despite not performing as well as ARGweaver in our evaluation criteria, Relate seems sufficient for comparisons of average trees across different regions in the genome.

Additionally, we showed that Relate’s inferences generally improve with sample size (Figures 6, 7, 8). This is expected from inference using the Li and Stephens (2003) copy algorithm, which tends to better approximate the genealogical process with larger samples sizes (Hubisz *et al*. 2020). Because Relate is fast enough, even for thousands of samples, it is preferred for large numbers of genomes - not only because ARGweaver is not scalable for such large sample sizes but also because Relate inference tends to improve with larger sample sizes (Hubisz and Siepel 2020).

The framework of tsinfer and tsdate is also based on the Li and Stephens (2003) model, and it additionally takes advantage of the succinct tree sequence data structure that makes it scalable to even larger sample sizes than Relate, and at least an order of magnitude larger than tested here (Wohns *et al*. 2022). Although we did not find an improvement of tsinfer+tsdate estimates with increasing sample sizes in the range we tested (4 to 200 haplotypes), our analyses cannot rule out the possibility of better tsinfer+tsdate inference at larger sample sizes.

Increasing the mutation to recombination ratio in simulations improved point estimates from ARGweaver but did not improve posterior calibration (Figure 5). This lack of improvement of the posterior sampling can be explained by lack of convergence and could potentially be improved by increasing the number of MCMC iterations. Although the statistics recorded by ARG-weaver at each iteration (likelihood, number of recombinations, etc.) show convergence (Figure S9, Table S1), we observed that certain pairwise coalescence times did not converge in the simulations with increased mutation to recombination ratio (Table S2 see more discussion in ARGweaver in Supplementary Materials).

### Limitations of our analyses and future directions

The focus of this study is the inference of coalescence times under the standard neutral coalescent, assuming all parameter values of this model are known and correctly provided to the programs performing inference. In other words, our goal was to investigate the performance of the ARG inference methods when the underlying assumptions are met. We have not explored how the methods perform under more complex demographic models and in the presence of natural selection, when the underlying assumptions are not met, but this is clearly an important future direction.

We also restrict our analyses to small sample sizes relative to what is possible for Relate and tsinfer+tsdate. However, increasing sample sizes up to 200 samples does not consistently improve performances of these methods. We also note that interesting discoveries have been made by applying ARG-based methods with similarly small sample sizes, *e*.*g*. Hubisz *et al*. (2020) analysed gene flow between archaic and modern humans using five genomes: two Neanderthals, one Denisovan and two modern humans.

Other factors not explored here could also be relevant for applications to real data. For example, sequencing or phasing errors could reduce the performance of all methods. Each of the methods compared here deal with these problems in a different way. Both Relate and tsinfer require phased data. While Speidel *et al*. (2019) argue that Relate is robust to errors in computational phasing, Kelleher *et al*. (2019) acknowledge that phasing errors could reduce the performance of tsinfer. ARGweaver is the only method of the three that supports unphased data, by integrating over all possible phases. However, the performance of the program on unphased data has not been evaluated in this study.

Relate takes sequencing errors into account by allowing some mutations that are incompatible with the tree topology in its tree building algorithm. Some robustness to error is shown in Speidel *et al*. (2019, Figure S3). Tsdate also uses heuristics in the ancestral haplotype reconstruction stage to increase its robustness to genotyping errors (Kelleher et al. 2019), and its newest version also accounts for recurrent mutation. ARGweaver can deal with genotyping errors statistically, using genotype likelihoods and integrating over all possible genotypes (Hubisz and Siepel 2020). In addition, it can take into account local variation in coverage and mapping quality, all of which are features not tested here. ARGweaver can also incorporate a map of variable mutation rates. ARGweaver, Relate and tsinfer can all incorporate maps of variable recombination rates across the genome, a feature which was not used in our constant rate simulations.

In our standard simulations, we use mutation rate equal to recombination rate, which is believed to be approximately true for humans. In reality, even if average recombination and mutation rates are similar, the average recombination rate is not distributed equally along the genome in humans and other mammals but is concentrated in recombination hotspots. Therefore, it is possible that ARG inference could be more accurate with real data, since local trees could span longer sequences separated by recombination hotspots.

### Recommendations for usage

Given that ARGweaver provides the most accurate coalescence times estimates and the most well-calibrated samples from the posterior distribution of coalescence times, we recommend using it whenever computationally feasible. However, it is highly computationally demanding and its usage can become unfeasible with sample sizes close to 100. Running ARGweaver on small segments of sequence (5Mb or 250kb Fig. S8, S7) gave similar results to applications on 100Mb segments, making the program highly parallelizable, at least for the purpose of estimating pairwise coalescence times.

When ARGweaver is computationally prohibitive, Relate and tsinfer+tsdate are viable alternative options. However, we emphasize that we have only examined coalescence time estimates, and for other downstream uses of ARG inference that do not rely mostly on coalescence times, the tradeoffs between these methods could be different. See Deng *et al*. (2021) for a comparison of these methods in the context of estimating recombination rates.

## Data availability

The data underlying this article are available in GitHub https://github.com/deboraycb/ARGsims.

## Acknowledgements

We would like to thank Leo Speidel for identifying an error in the extraction of coalescence times from the Relate output in a previous version of the manuscript. We also thank Nicholas Barton, Adam Siepel, Ziyi Mo, Jerome Kelleher, Yan Wong, Wilder Wohns and other reviewers for constructive and detailed reviews that greatly improved this paper.

## Funding

This material is based upon work supported by NSF Graduate Research Fellowship No. 2146752 awarded to Andrew Vaughn and by NIH grant R01GM138634 awarded to Rasmus Nielsen.

## Supplementary Materials

### Abstract in Portuguese - Resumo em português

The Portuguese translation of the abstract was done using Google Translate and corrected by Débora Y. C. Brandt, who is a native Portuguese speaker.

A tradução do resumo deste artigo para o português foi feita usando o Google Tradutor, seguida de correção manual por Débora Y.

C. Brandt, que tem o português como língua materna.

O grafo de recombinação ancestral (GRA, ou ARG na sigla em inglês) é uma estrutura que descreve o conjunto de genealogias locais ao longo do genoma, para um conjunto de sequências de DNA amostradas. Métodos computacionais desenvolvidos recentemente geraram um progresso impressionante na possibilidade de estimar genealogias de todo o genoma para um grande número de amostras. Além de inferir um único ARG, alguns desses métodos também podem fornecer diversos ARG amostrados de uma distribuição *a posteriori*. Obter uma boa amostra de ARGs é crucial para quantificar a incerteza estatística e para estimar parâmetros populacionais, como tamanho efetivo da população, taxa de mutação e idade de alelos. Neste trabalho, usamos simulações sob o modelo coalescente neutro padrão para comparar as estimativas de tempos de coalescência par-a-par de três programas amplamente utilizados para a inferência de ARGs: ARGweaver, Relate e tsinfer+tsdate. Comparamos 1) os tempos de coalescência simulados com os tempos inferidos em cada *locus*; 2) a distribuição dos tempos de coalescência par-a-par para todos os *loci* com a distribuição exponencial que seria esperada; 3) se os tempos de coalescência amostrados possuem as propriedades esperadas de uma distribuição *a posteriori* bem calibrada. Descobrimos que os tempos de coalescência inferidos locus-a-locus pelo programa ARGweaver são os mais precisos, e que geralmente os tempos de coalescência inferidos pelo programa Relate são mais precisos do que os inferidos por tsinfer+tsdate. No entanto, os três métodos tendem a superestimar tempos de coalescência baixos e subestimar os altos. Por fim, as amostras da distribuição *a posteriori* geradas pelo programa ARGweaver refletem uma distribuição mais próxima da distribuição *a posteriori* esperada do que as amostras geradas pelo programa Relate, mas essa precisão mais alta é acompanhada de custo computacional muito mais elevado. Portanto, a escolha do melhor método a ser usado depende do número e comprimento das sequências amostradas, e do objetivo das análises em que se deseja usar o ARG. Por fim, oferecemos algumas recomendações de uso desses métodos para diferentes fins.

#### Palavras-chave

Grafo de recombinação ancestral; ARGweaver; Relate; tsinfer; tsdate; simulação; calibração; distribuição *a posteriori*

### Figures

**Figure S1.**
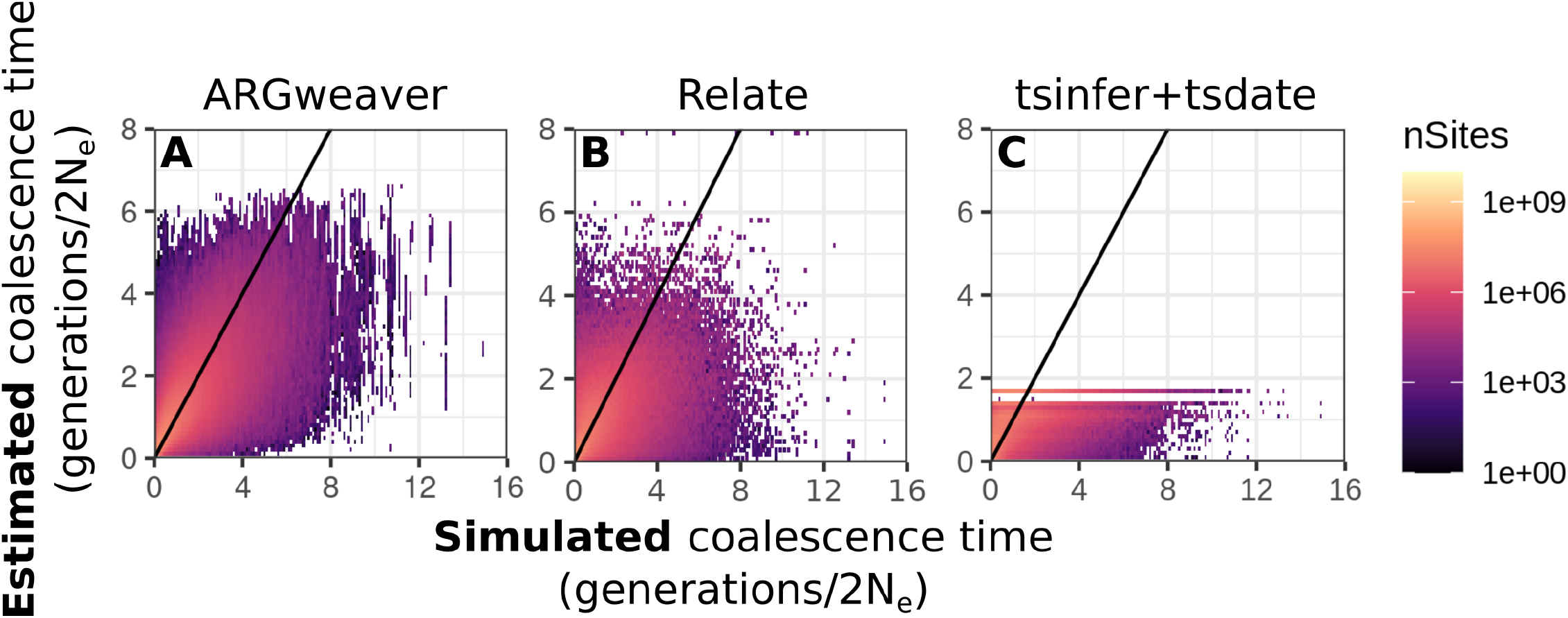
True pairwise coalescence time from msprime simulations compared to inferred coalescence time from (A) ARGweaver (B) Relate (C) tsdate. Note that axes are in linear scale. See Figure 3A, D, G for this data plotted on a logarithmic scale. These results are for simulations with n=8 samples (haplotypes), mutation and recombination rates of 2 *×* 10^−8^. Diagonal line shows x=y, points show the mean inferred coalescence time within a true coalescence time bin.

**Figure S2.**
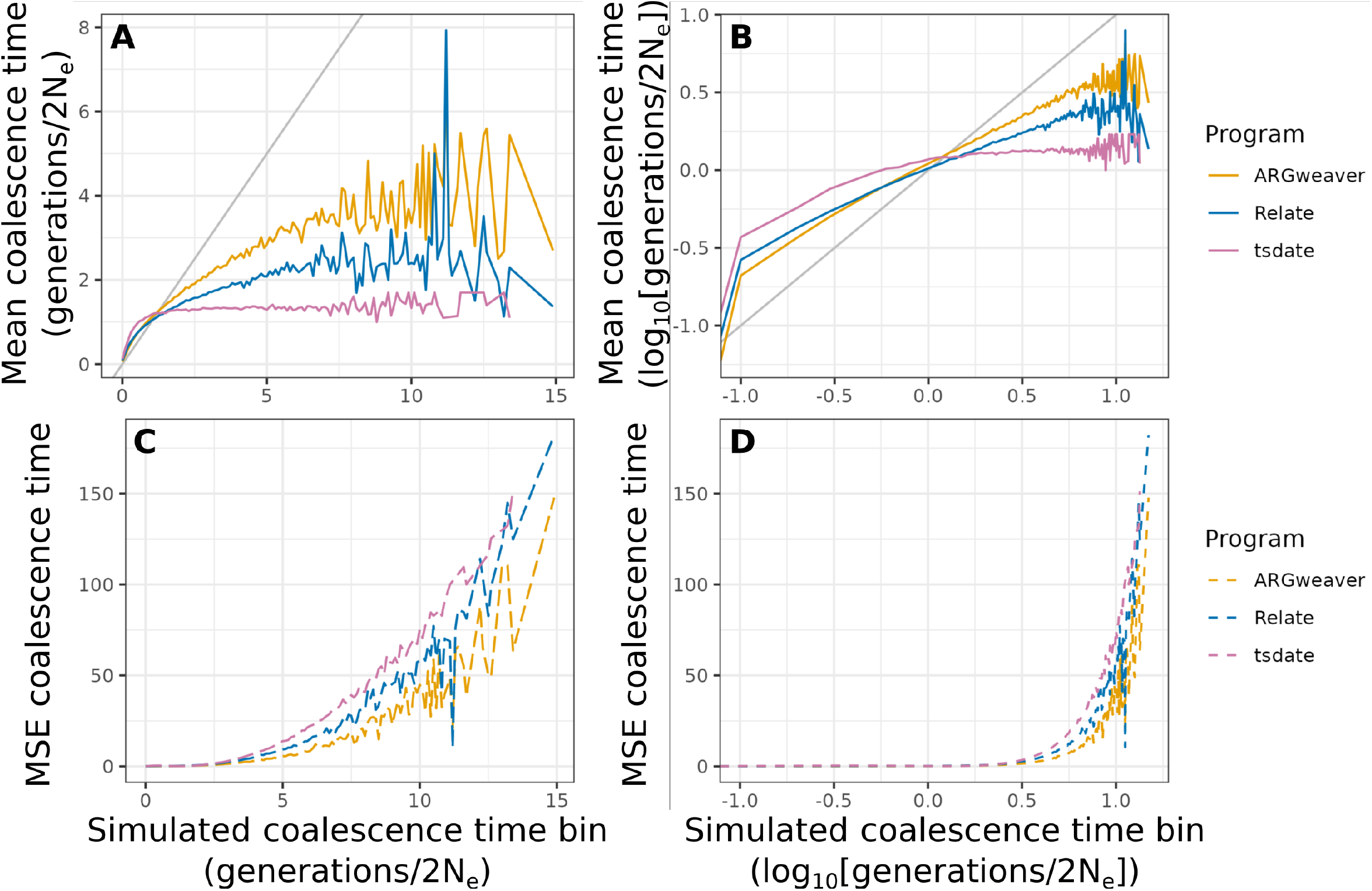
Mean (A,B) and mean squared error (C,D) of point estimates of pairwise coalescence times by ARGweaver, Relate and tsdate in each bin of size 0.1 of simulated coalescence times. Diagonal gray line in plots A and B show 1:1 line. These results are for simulations with n=8 samples, mutation and recombination rates of 2 *×* 10^−8^. Plots B and D are in log scale to highlight small values of coalescence times, which are the most abundant. Note that estimates are best (*i*.*e*. means in plots a and b are closer to the simulated value) at values near the expected mean coalescence time under the coalescent *(i*.*e*. 1 in the coalescent units of 2*N*_*e*_ generations).

**Figure S3.**
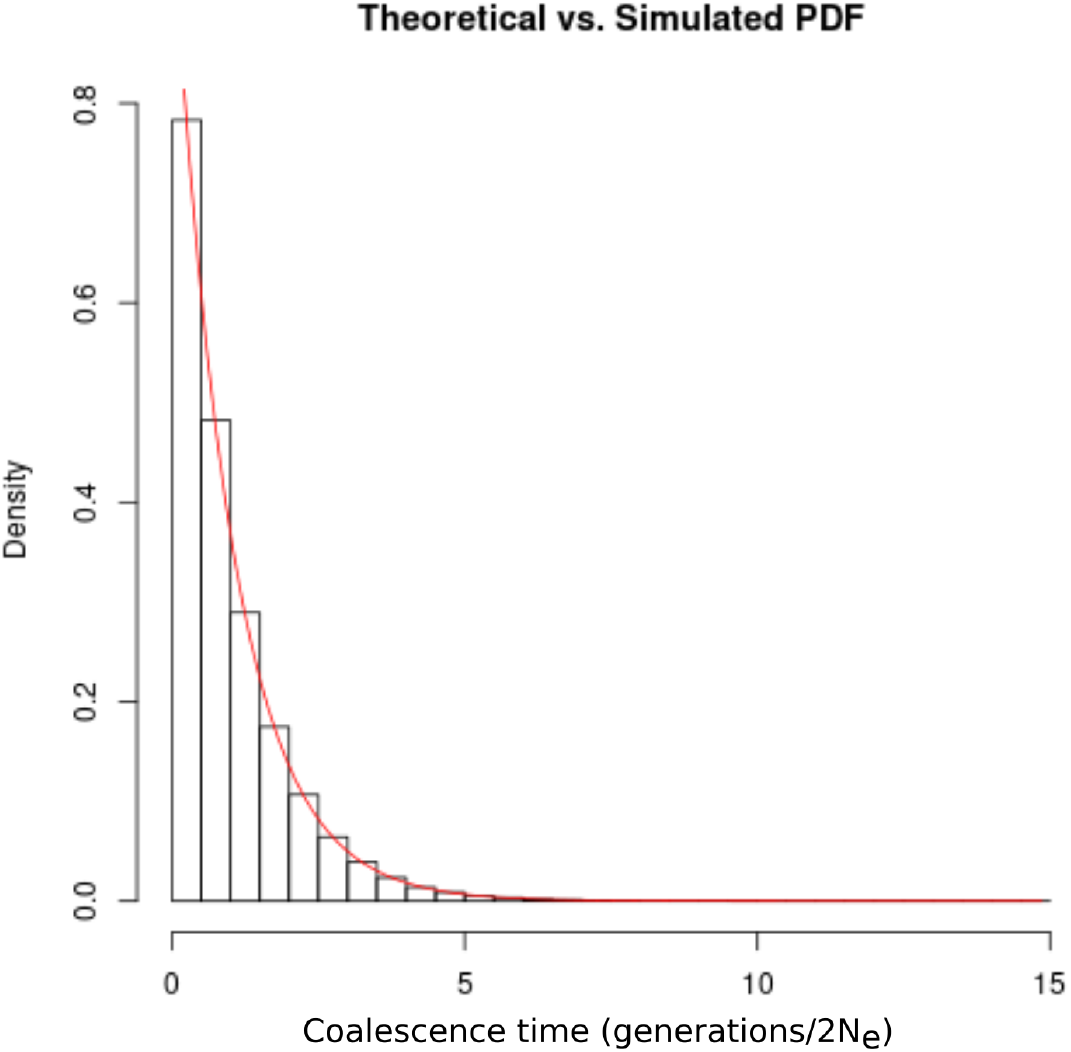
Histogram of the distribution of coalescence times in msprime simulations. Red line show expected exponential distribution with rate 1.

**Figure S4.**
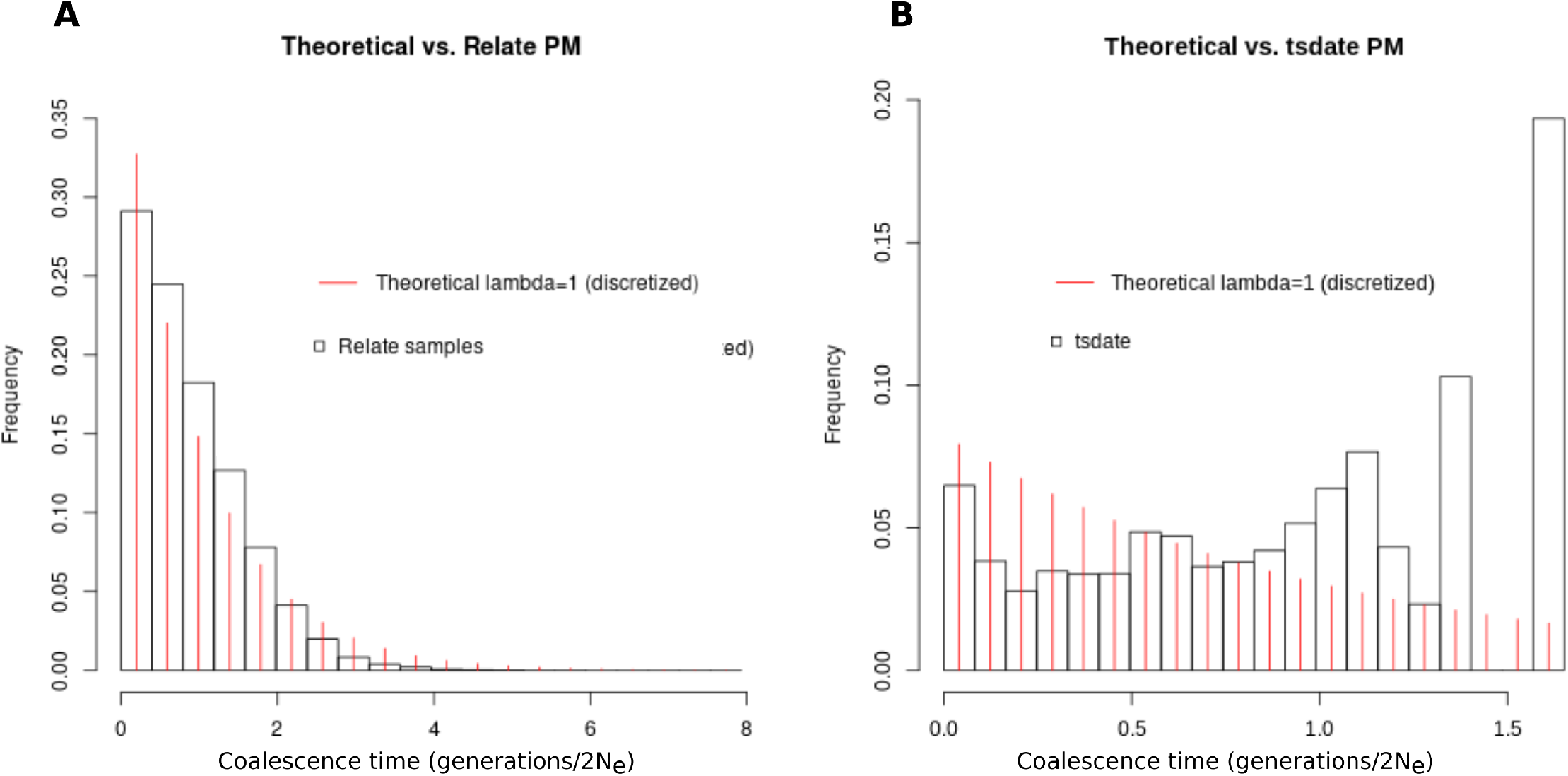
Distributions of pairwise coalescence times in Relate and tsdate without ARGweaver time discretization. These results are for simulations with n=8 samples, mutation and recombination rates of 2 *×* 10^−8^. (a) Relate, (b) tsdate, both with 20 equal size bins

**Figure S5.**
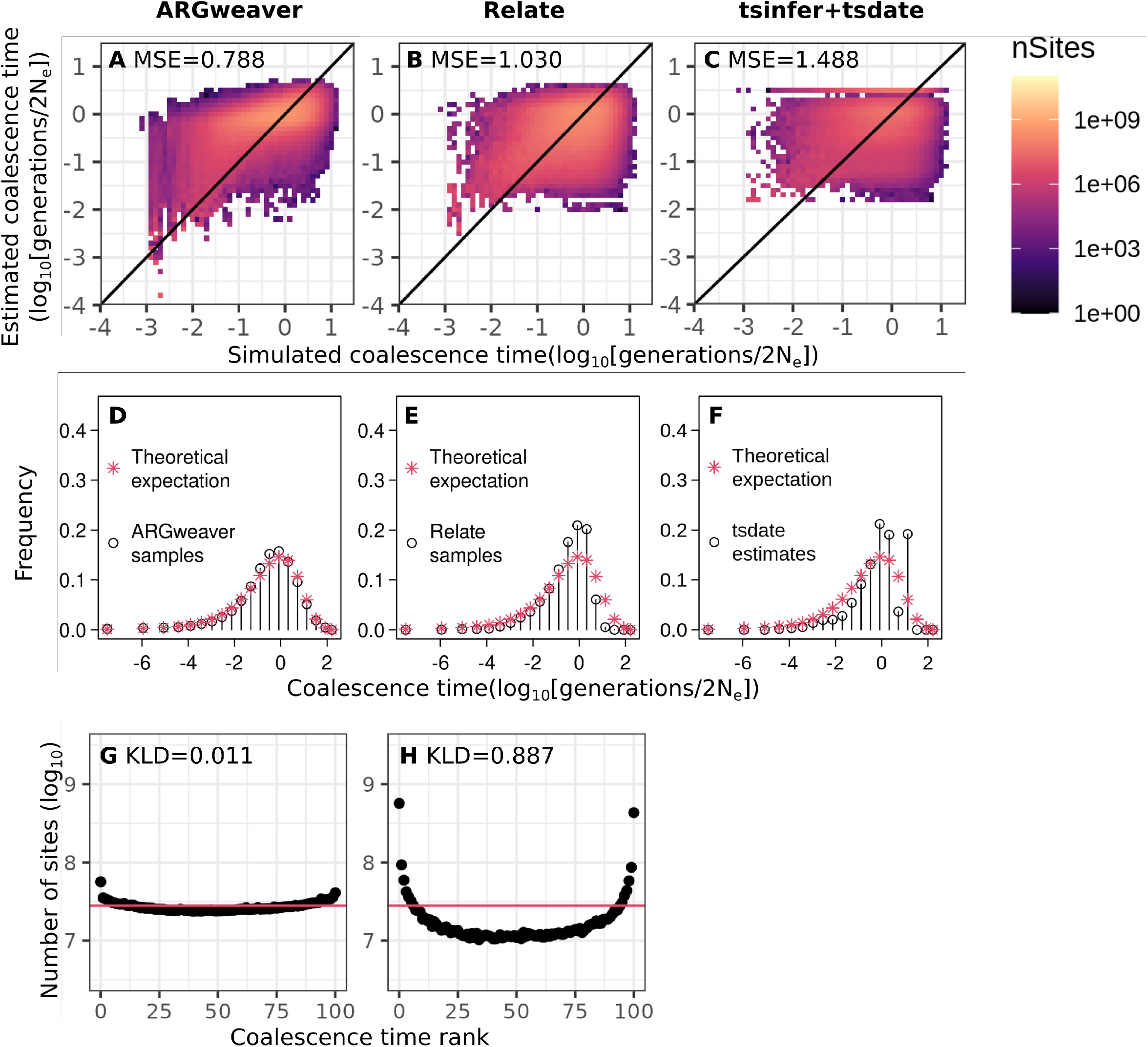
Point estimates (A-C), distribution of coalescence times (D-F) and counts of ranks from simulation-based calibration (G,H) from ARGweaver (A,D,C), Relate (B,E,H) and tsinfer+tsdate (C,F). Simulations with **reduced mutation rate** (*µ* = 2 *×* 10^−9^ and *ρ* = 2 *×* 10^−8^). Compared to simulations with mutation rate equal to recombination rate, mean square error (MSE) values are all larger (Figure 3), distributions of coalescence times deviate more from the theoretical expectation (Figure 4), and KLD is lower in ARGweaver but higher in Relate (Figure 5).

**Figure S6.**
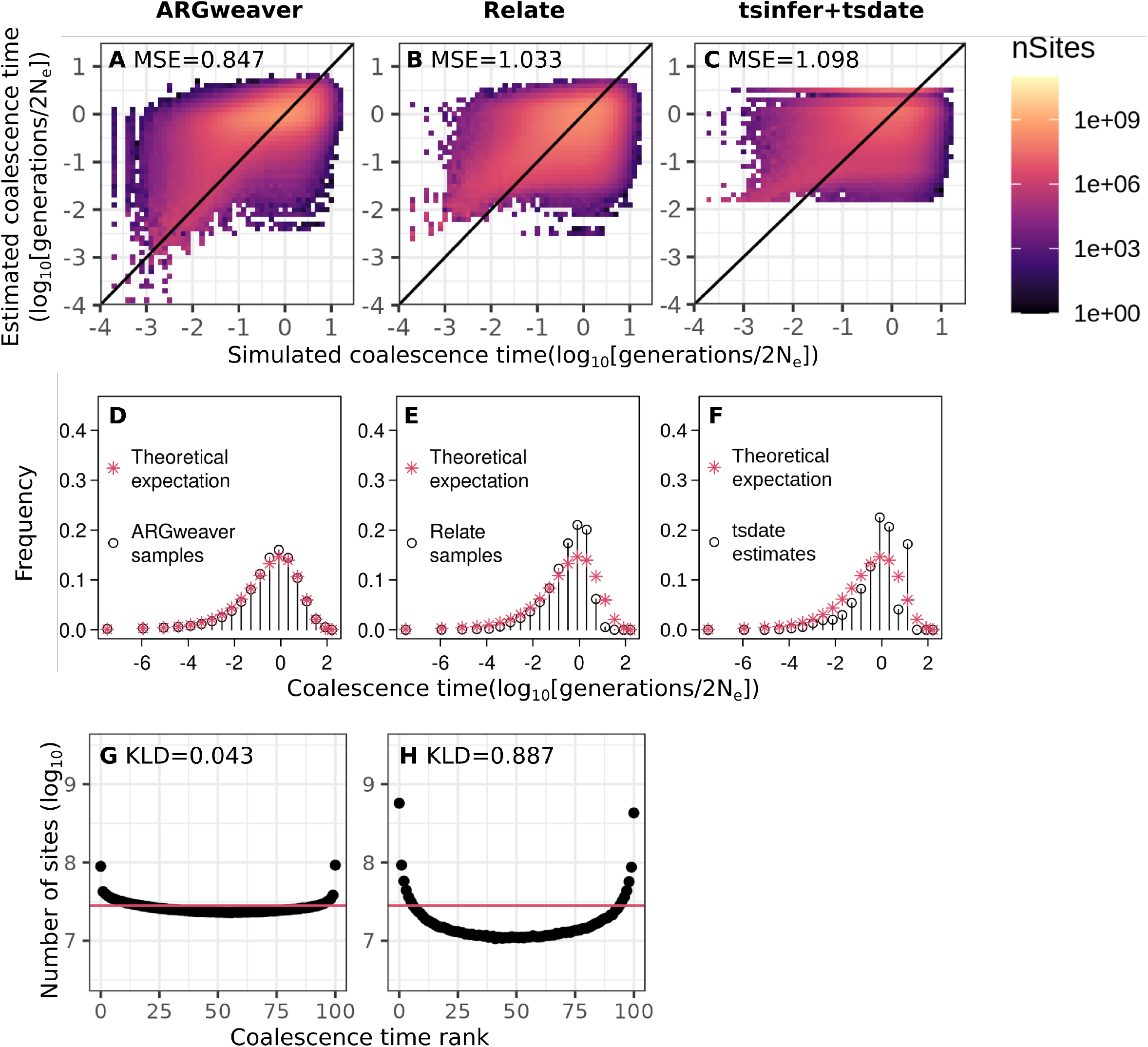
Point estimates (A-C), distribution of coalescence times (D-F) and counts of ranks from simulation-based calibration (G,H) from ARGweaver (A,D,C), Relate (B,E,H) and tsinfer+tsdate (C,F). Simulations with **increased recombination rate** (*µ* = 2 *×* 10^−8^ and *ρ* = 2 *×* 10^−7^). Compared to simulations with mutation rate equal to recombination rate, Mean square error (MSE) values are all larger (Figure 3), distributions of coalescence times deviate more from the theoretical expectation (Figure 4), and KLD is lower in ARGweaver, but higher in Relate (Figure 5).

**Figure S7.**
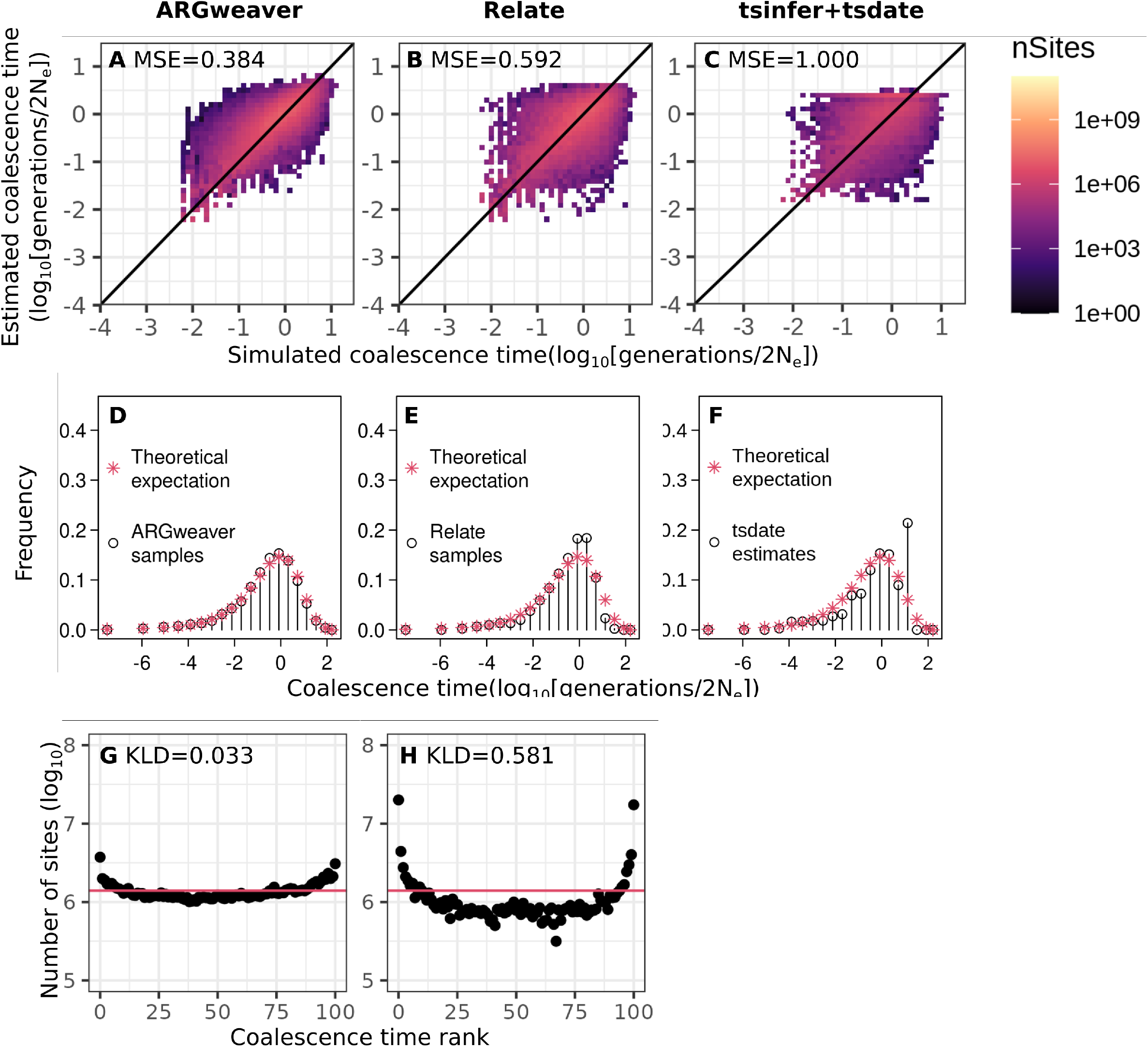
Point estimates (A-C), distribution of coalescence times (D-F) and counts of ranks from simulation-based calibration (G,H) from ARGweaver (A,D,C), Relate (B,E,H) and tsinfer+tsdate (C,F). Simulations with sample size of 8 haplotypes, *µ* = *ρ* = 2 *×* 10^−8^, and **input sequence length of 5Mb**.

**Figure S8.**
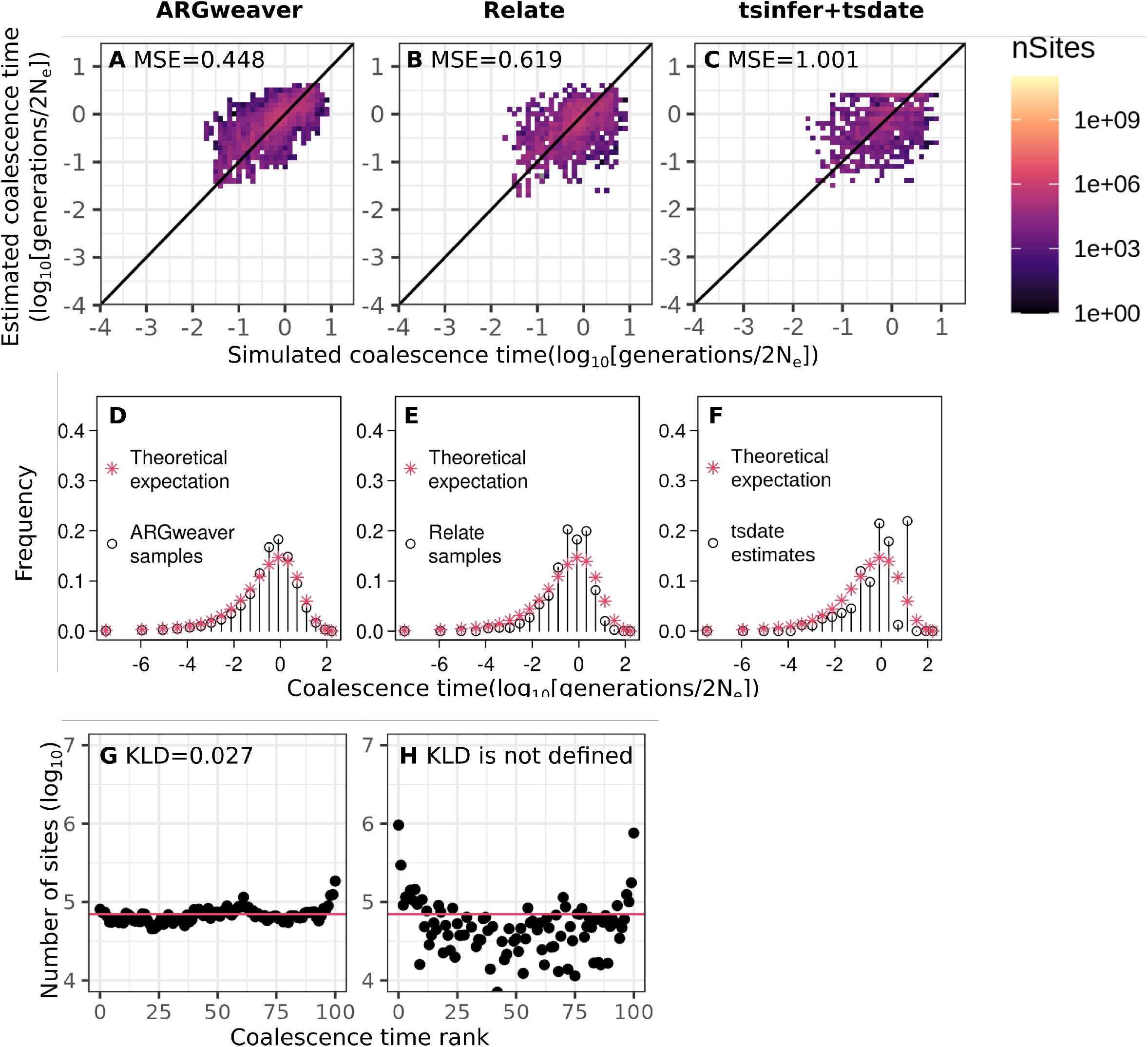
Point estimates (A-C), distribution of coalescence times (D-F) and counts of ranks from simulation-based calibration (G,H) from ARGweaver (A,D,C), Relate (B,E,H) and tsinfer+tsdate (C,F). Simulations with sample size of 8 haplotypes, *µ* = *ρ* = 2 *×* 10^−8^, and **input sequence length of 250kb**. In H, KLD is not defined because counts for one of the ranks is zero.

**Figure S9.**
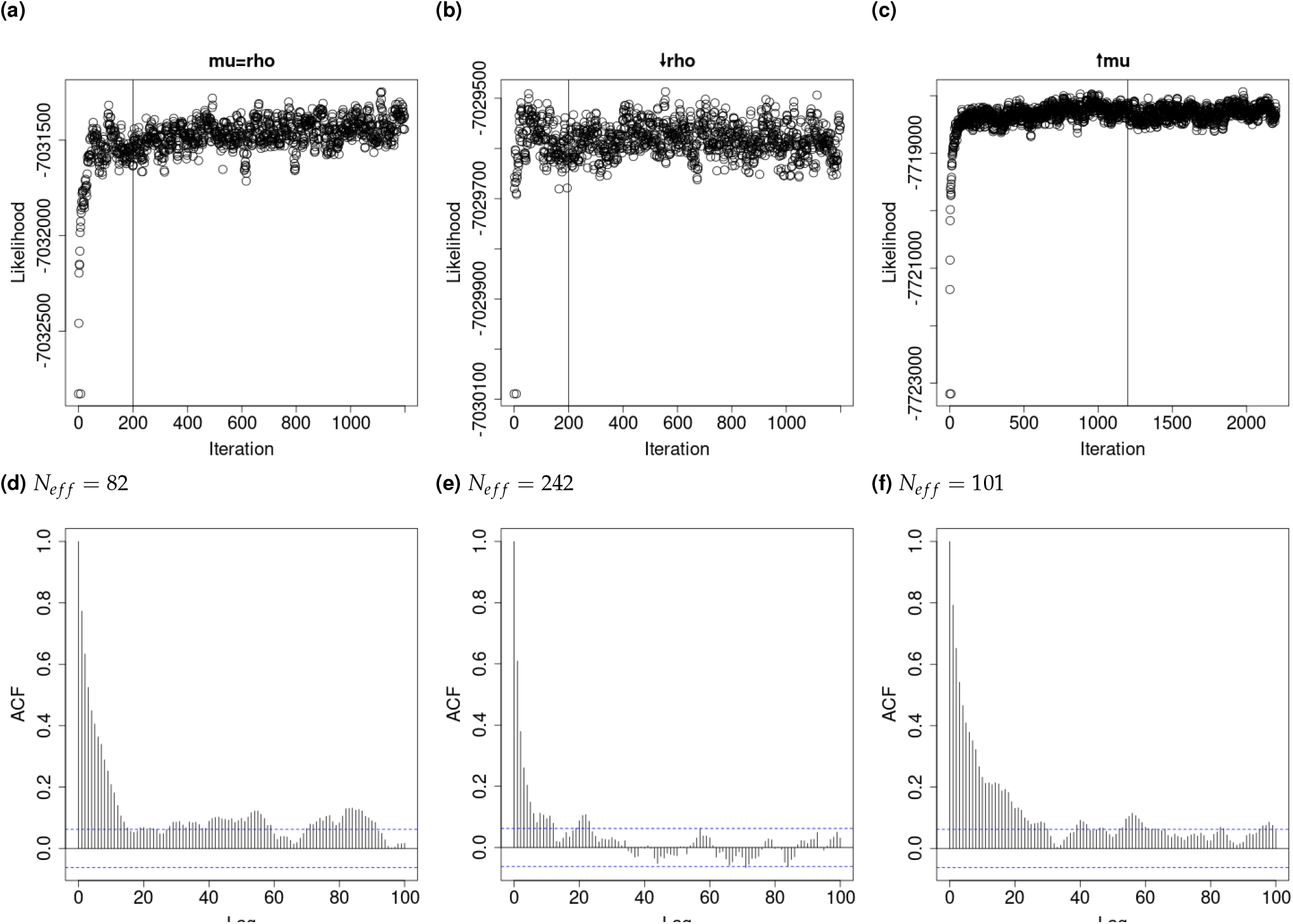
ARGweaver likelihood traces (top) and autocorrelation between consecutive MCMC iterations (bottom, also showing effective sample sizes (*N*_*e f f*_)) for the number of iterations used in the main text. Left column: simulations with 8 haplotypes, mutation rate equal to the recombination rate(2 *×* 10^−8^). Potential scale reduction factor (PSRF) is 1.02, upper confidence interval (CI) is 1.05. Middle column: simulations with recombination rate decreased to 2 *×* 10^−9^. PSRF is 1.04, upper CI is 1.11. For both of these simulated datasets we used a burn in of 200 iterations (indicated by vertical line) and ran them for 1200 iterations in total, sampling every 10th iteration. Right column: simulations with mutation rate increased to 2 *×* 10^−7^. PSRF is 1.01, upper CI is 1.02. For this dataset we used a burn in of 1200 iterations (indicated by vertical line) and ran them for 2200 iterations in total, sampling every 10th iteration.

**Figure S10.**
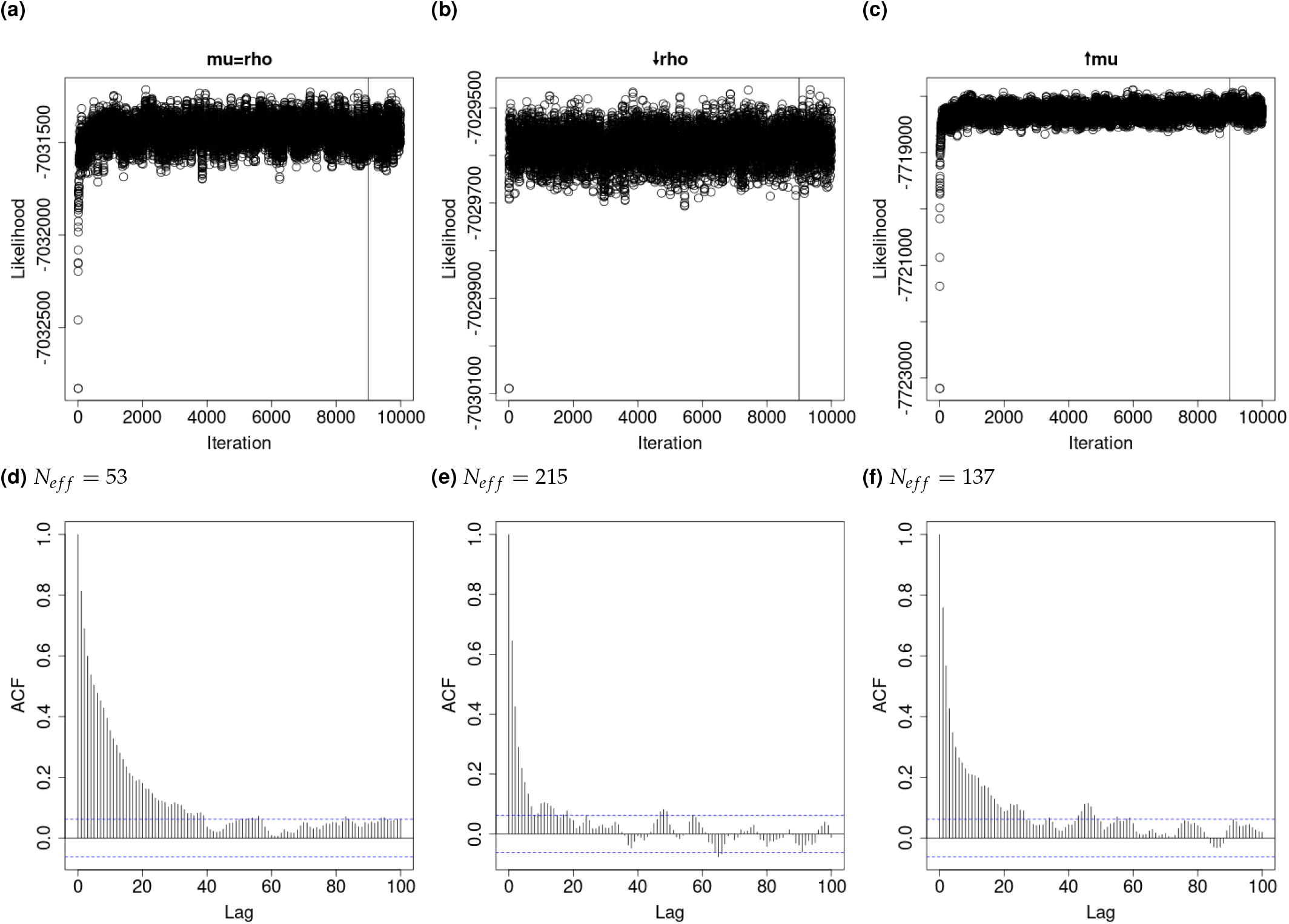
Similar to Figure S9, but running ARGweaver for 10 thousand iterations, with a burn in of 9 thousand applied before calculating effective sample sizes, to keep the same number of samples (1000). ARGweaver likelihood traces (A,B,C) and autocorrelation between consecutive MCMC iterations (D,E,F). Left column: simulations with 8 haplotypes, mutation rate equal to the recombination rate (2 *×* 10^−8^). Middle column: simulations with recombination rate decreased to 2 *×* 10^−9^. Right column: simulations with mutation rate increased to 2 *×* 10^−7^.

**Figure S11.**
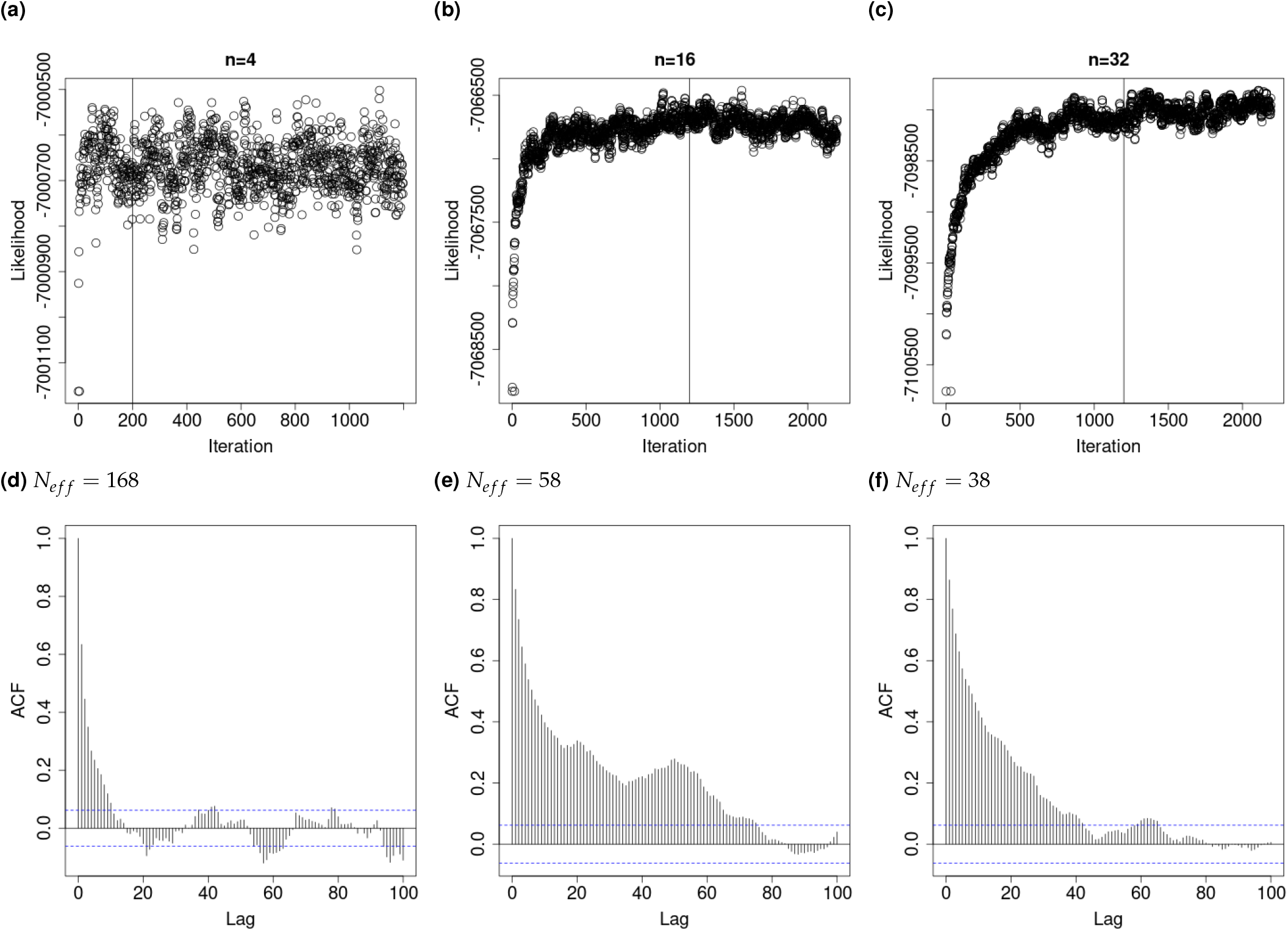
ARGweaver likelihood traces (top) and autocorrelation between consecutive MCMC iterations (bottom). A,D: simulations with 4 haplotypes, mutation rate equal to recombination rate (2 *×* 10^−8^). For this simulated dataset we used a burn in of 200 iterations (indicated by vertical line) and ran them for 1200 iterations in total, sampling every 10th iteration. B,E: simulations with 16 haplotypes. C,F: simulations with 32 haplotypes. For both of these datasets we used a burn in of 1200 iterations (indicated by vertical line) and ran them for 2200 iterations in total, sampling every 10th iteration.

**Figure S12.**
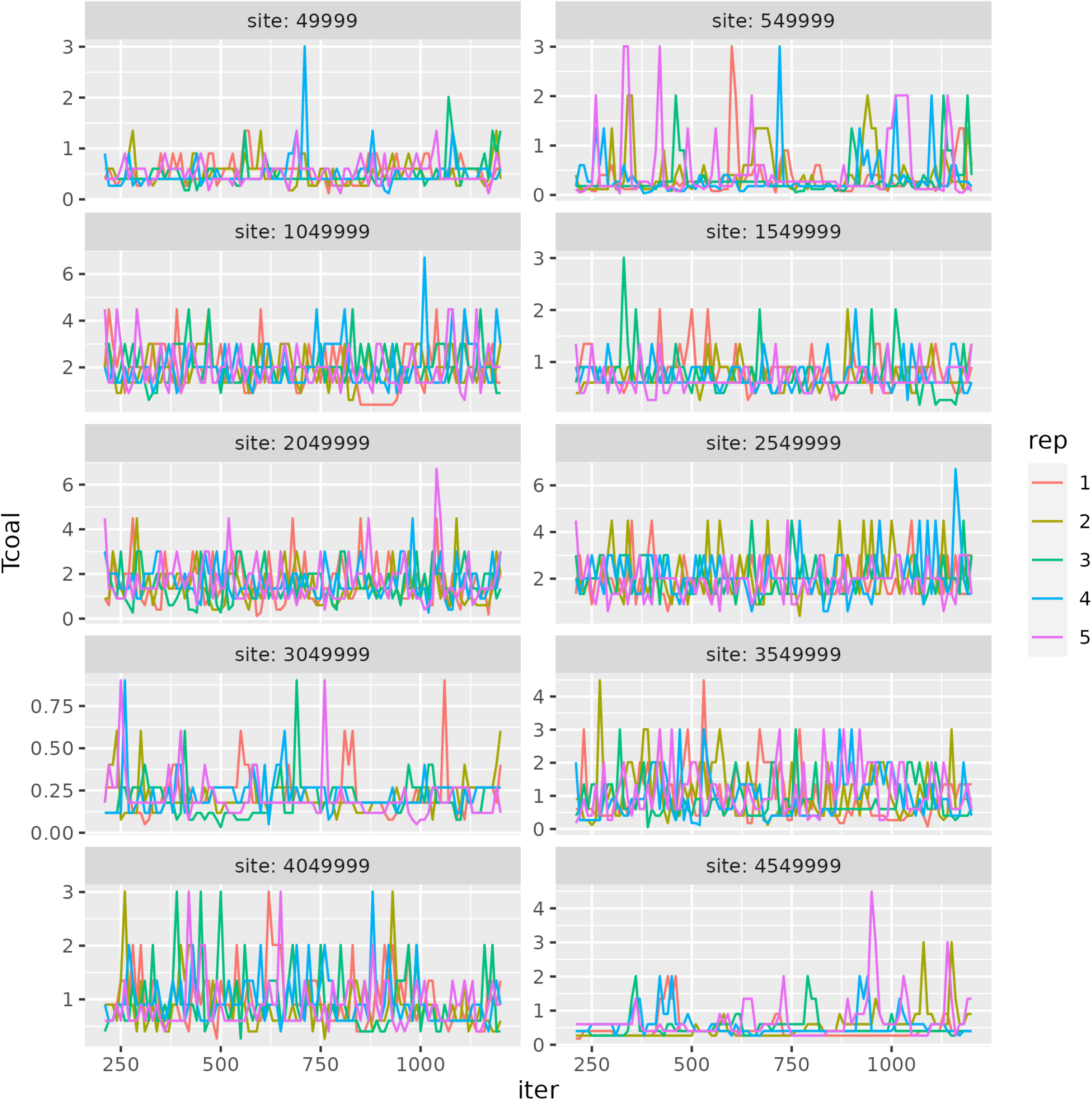
Coalescence times for one pair of samples inferred by 5 independent runs of ARGweaver at 10 sites equally spaced sites along the 5Mb sequence. Simulations with 8 samples and mutation rate equal to recombination rate.

**Figure S13.**
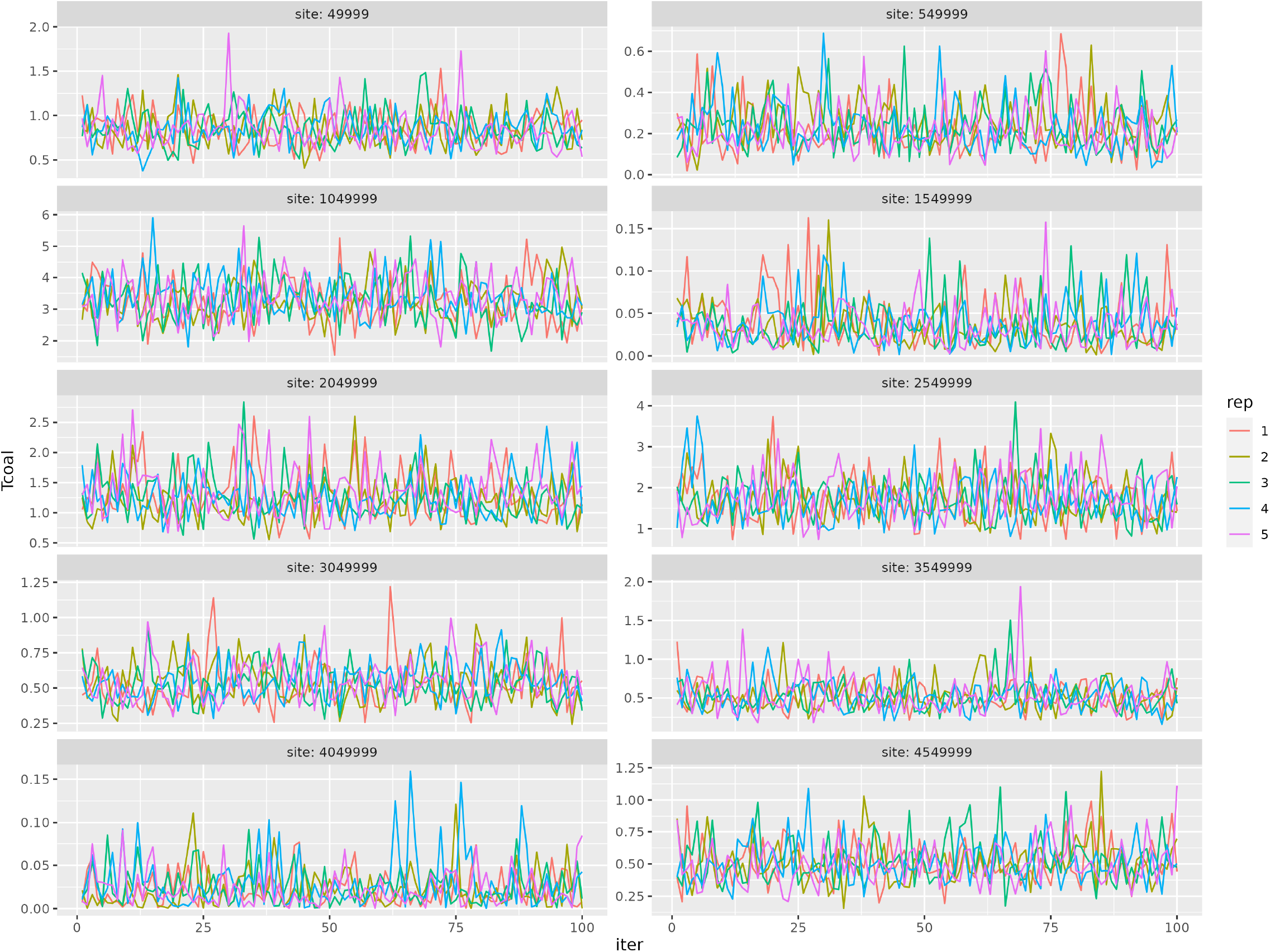
Coalescence times for one pair of samples inferred by 5 independent runs of Relate at 10 sites equally spaced sites along the 5Mb sequence. Simulations with 8 samples and mutation rate equal to recombination rate.

**Figure S14.**
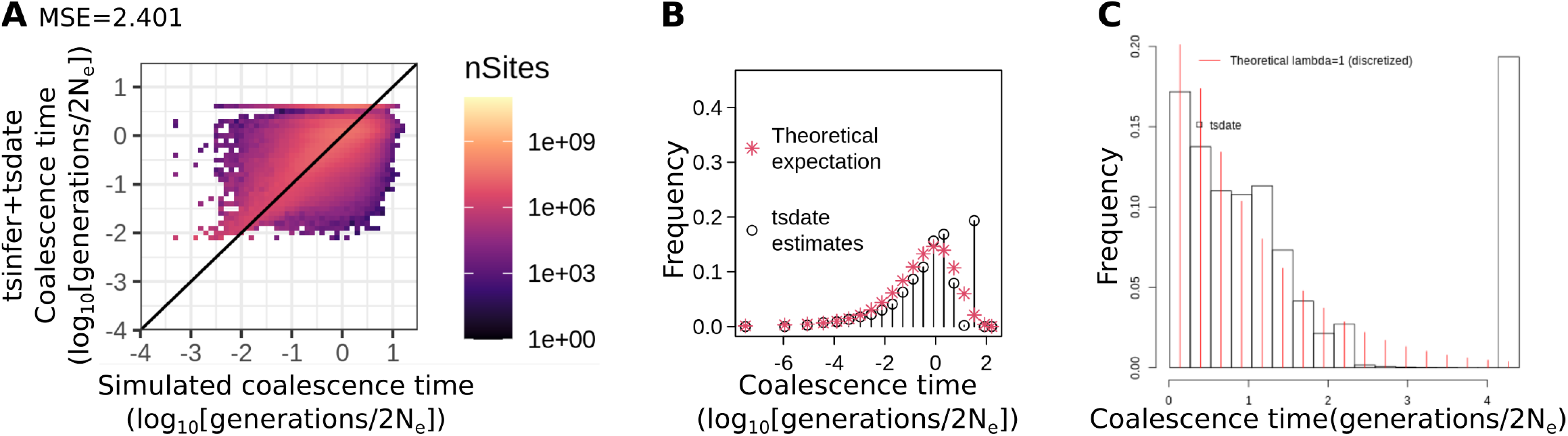
Tsdate results with a **prior grid constructed with timepoints=100**. (A) Comparisons of estimated and simulated point estimates of pairwise coalescence times. (B) Comparisons of the distribution of coalescence times to the expected exponential distribution, using ARGweaver time discretization bins. (C) Same as B, but without imposing ARGweaver time discretization.

**Figure S15.**
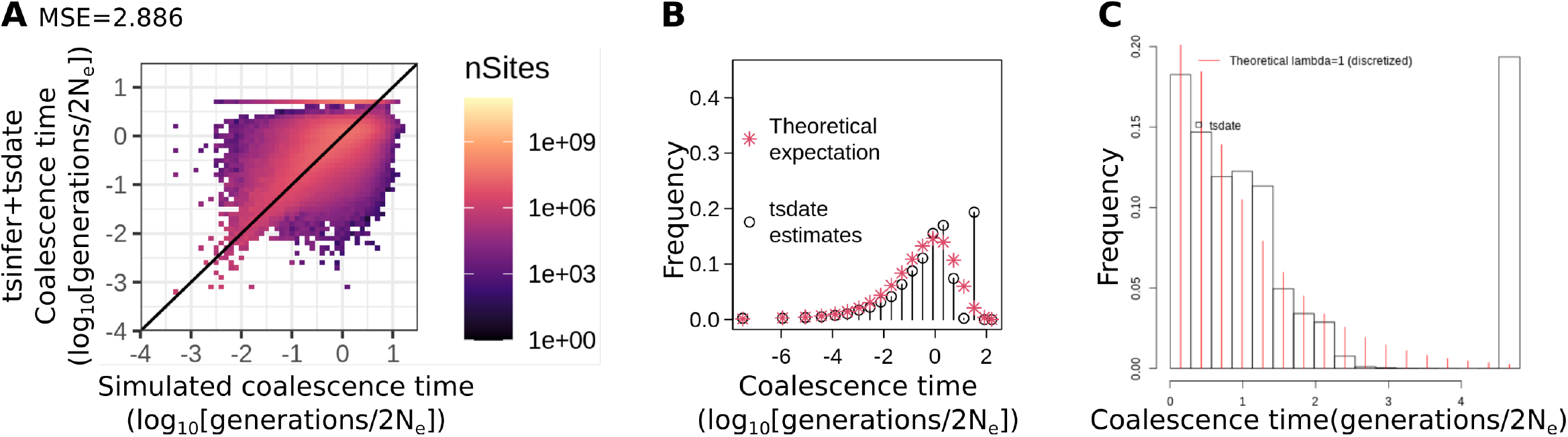
Tsdate results with a **prior grid constructed with a maximum value of 12**. (A) Comparisons of estimated and simulated point estimates of pairwise coalescence times. (B) Comparisons of the distribution of coalescence times to the expected exponential distribution, using ARGweaver time discretization bins. (C) Same as B, but without imposing ARGweaver time discretization.

**Figure S16.**
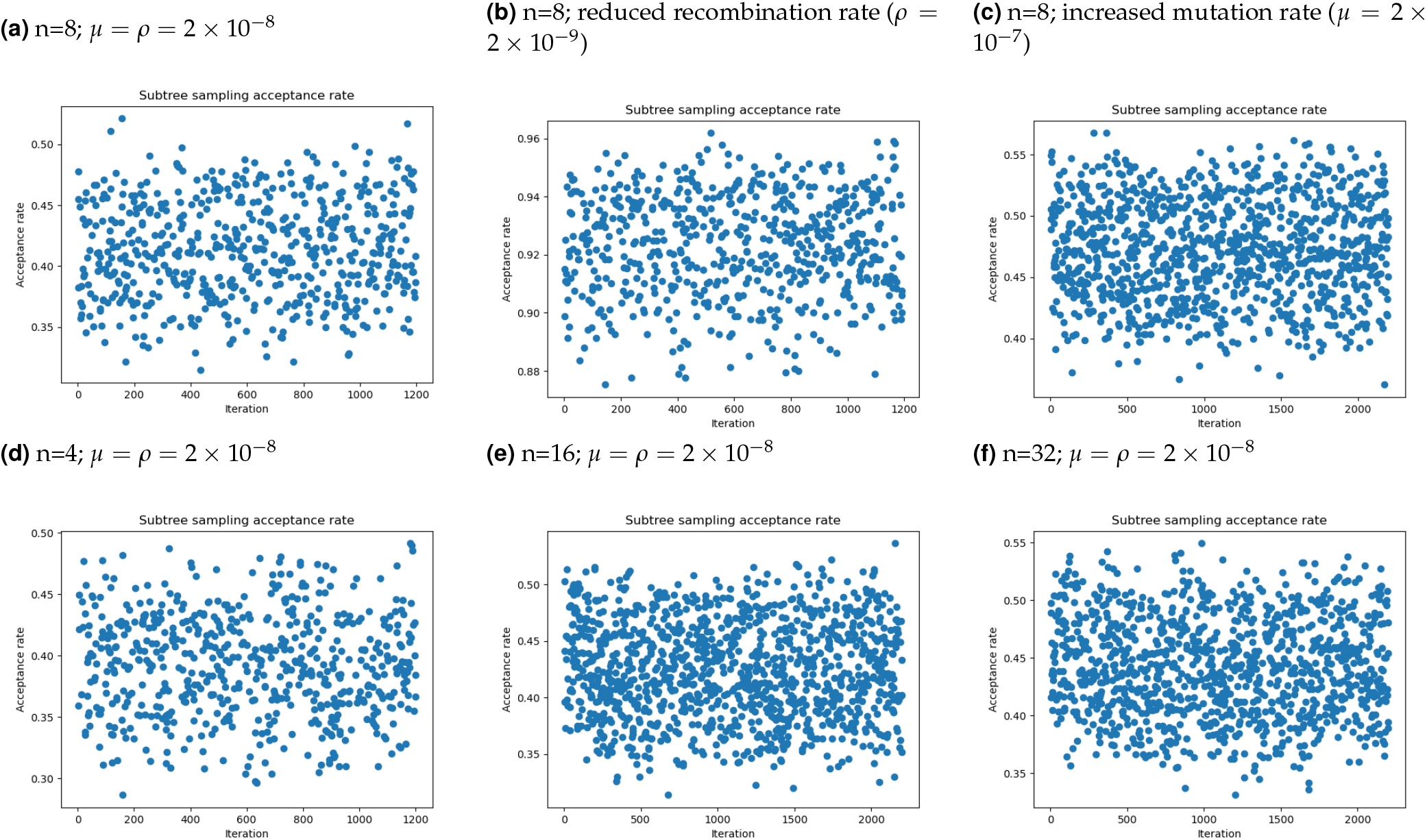
Acceptance rate from ARGweaver subtree sampling steps in one 5Mb region of each simulation.

**Figure S17.**
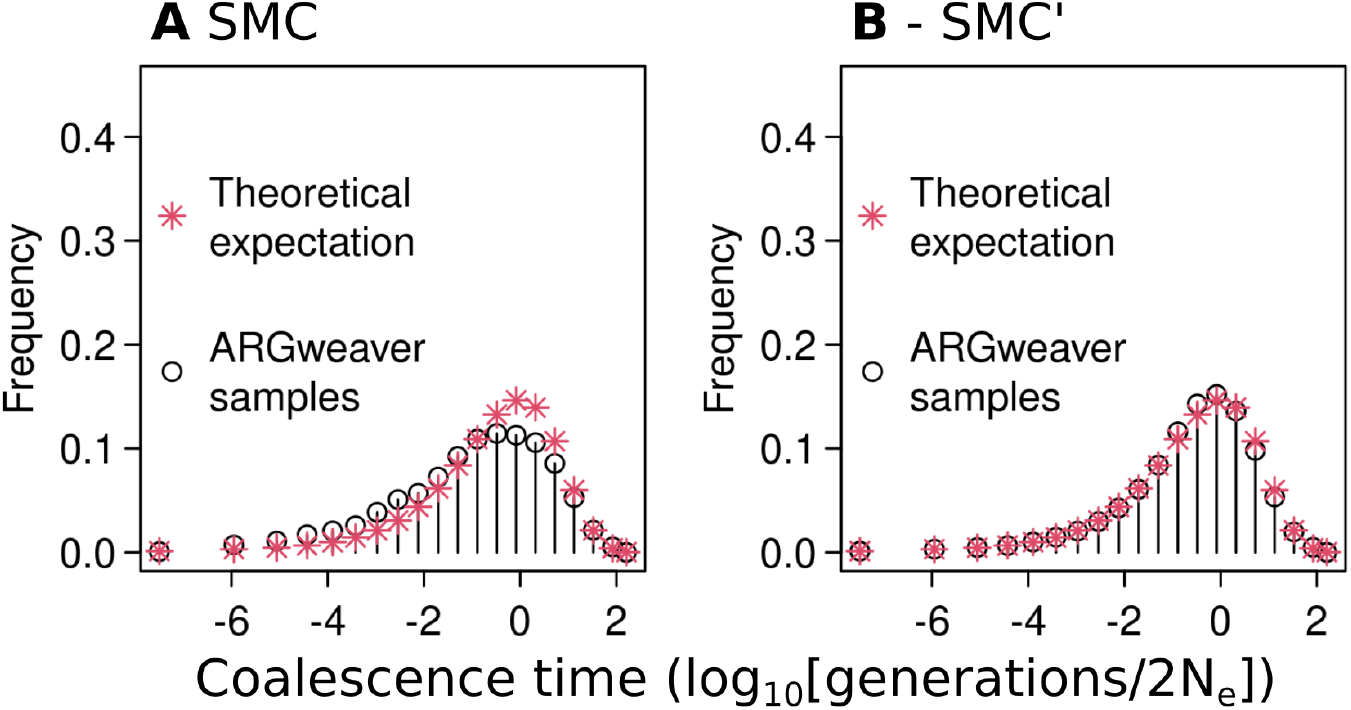
Distribution of coalescence times in msprime simulations using the SMC (A) or SMC’ model (B). ARGweaver inference is done using the same model used in the simulations.

**Figure S18.**
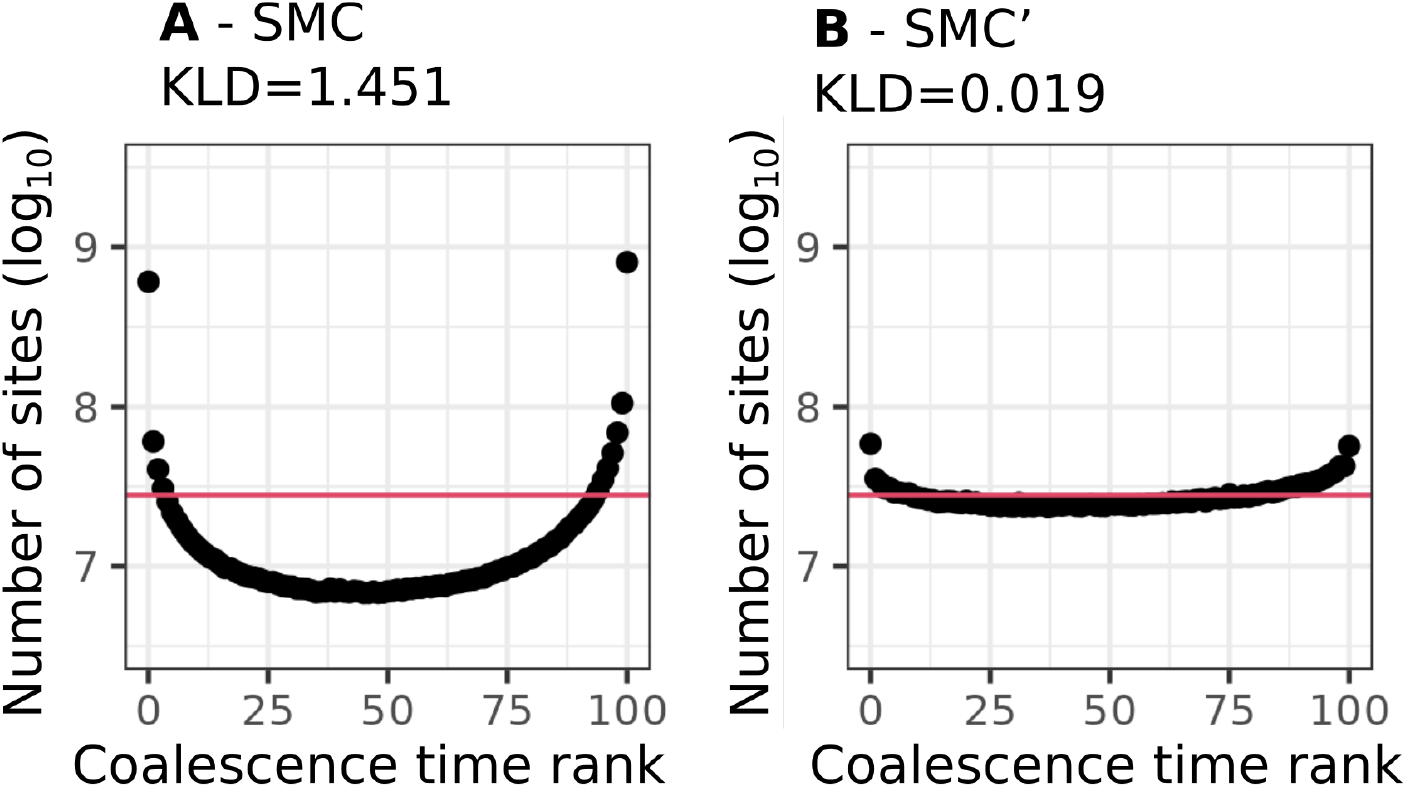
Simulation-based calibration results in msprime simulations using the SMC (A) or SMC’ model (B). ARGweaver inference is done using the same model used in the simulations.

**Figure S19.**
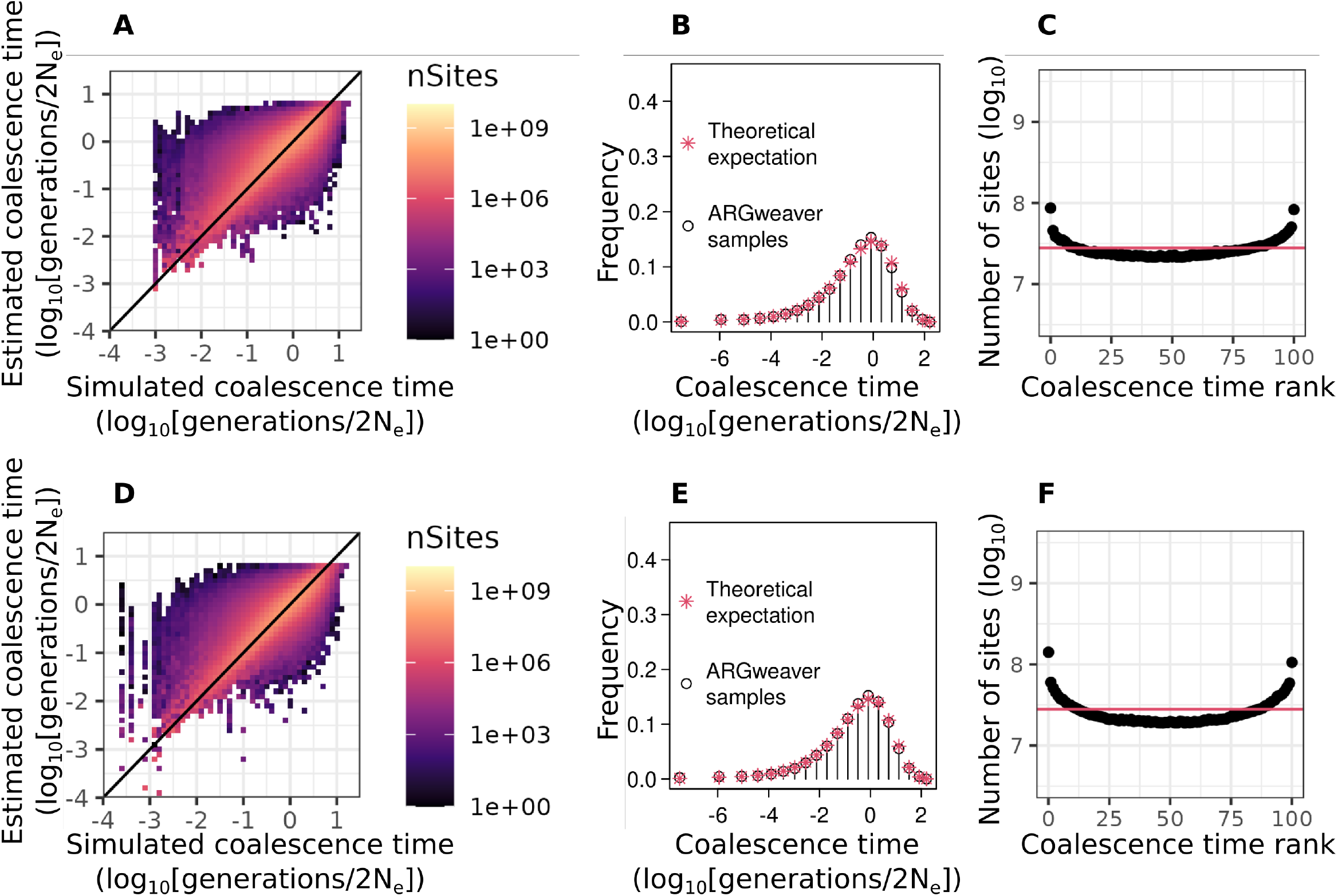
Evaluation of ARGweaver point estimates (A,D), distribution of coalescence times (B,E) and posterior calibration (C,F) for simulations with mutation rate to recombination rate ratio of 2 (A-C, *µ* = 4 *×* 10^−8^, *ρ* = 2 *×* 10^−8^) and mutation rate to recombination rate ratio of 4 (D-F, *µ* = 8 *×* 10^−8^, *ρ* = 2 *×* 10^−8^)

**Figure S20.**
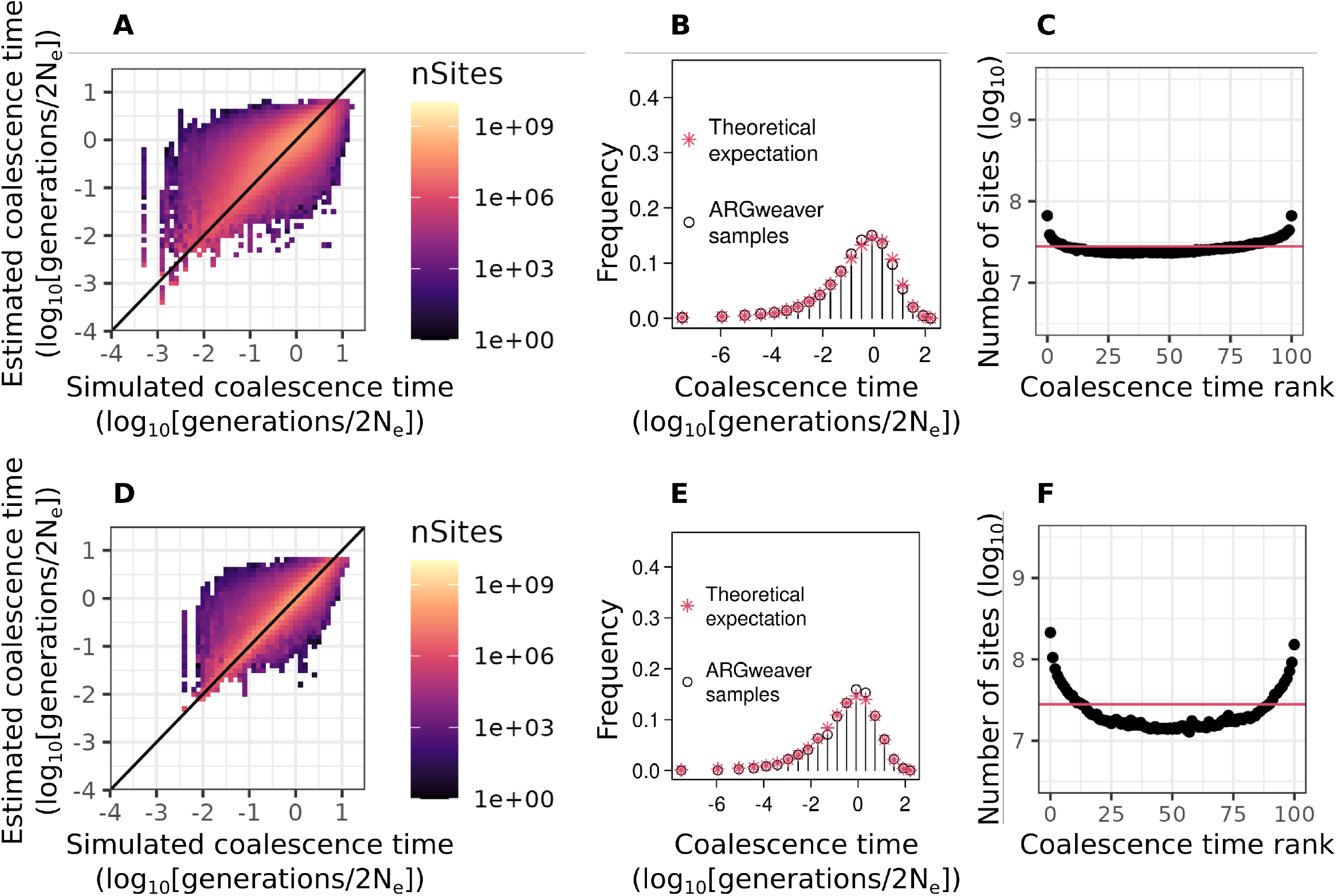
Evaluation of ARGweaver point estimates (A,D), distribution of coalescence times (B,E) and posterior calibration (C,F) with simulations under the Jukes and Cantor (1969) mutational model. A-C: simulations with 8 haplotypes and *µ* = *ρ* = 2 *×* 10^−8^. D-F: simulations with 8 haplotypes and *µ* = 2 *×* 10^−8^ and *ρ* = 2 *×* 10^−9^

**Figure S21.**
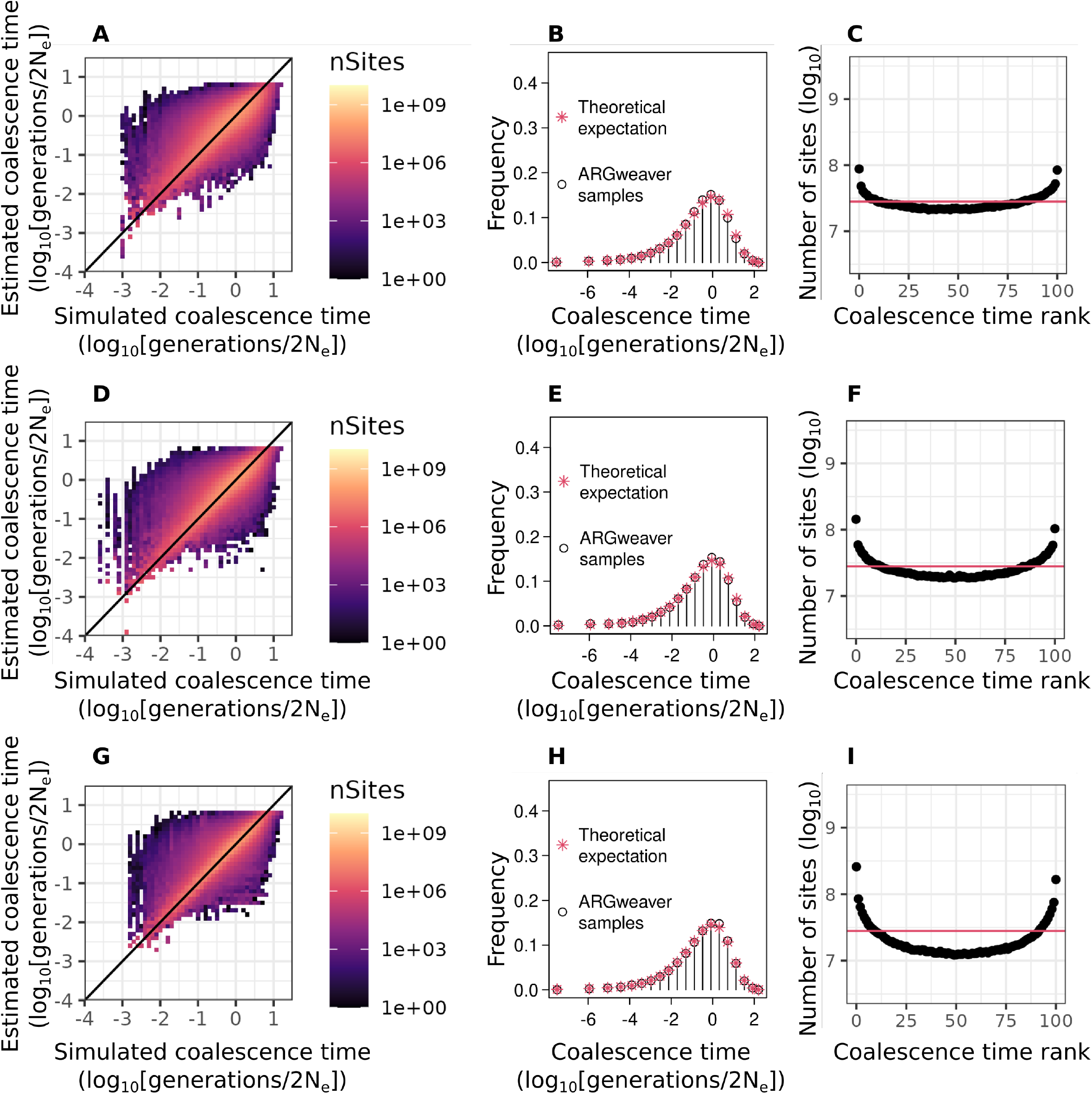
Evaluation of ARGweaver point estimates (A,D,G), distribution of coalescence times (B,E,H) and posterior calibration (C,F,I) with simulations under the Jukes and Cantor (1969) mutational model. In all cases we simulated 8 haplotypes and used *ρ* = 2 *×* 10^−8^. A-C: *µ* = 4 *×* 10^−8^. D-F: *µ* = 8 *×* 10^−8^. G-I: *µ* = 2 *×* 10^−7^

**Table S1.** Potential scale reduction factor point estimates (PSRF), their upper confidence intervals (C.I.) and effective sample sizes (*N*_*e f f*_) for ARGweaver stats. *µ*: mutation rate, *ρ*: recombination rate.

| | $\mu = \rho = 2 \times 10^{-8}$ | | | $\rho = 2 \times 10^{-9}$ | | | $\mu = 2 \times 10^{-7}$ | | |
| --- | --- | --- | --- | --- | --- | --- | --- | --- | --- |
| | PSRF | C.I. | $N_{eff}$ | PSRF | C.I. | $N_{eff}$ | PSRF | C.I. | $N_{eff}$ |
| prior | 1.06 | 1.15 | 224 | 1.01 | 1.03 | 494 | 1.00 | 1.01 | 216 |
| likelihood | 1.02 | 1.05 | 294 | 1.04 | 1.11 | 964 | 1.01 | 1.02 | 499 |
| joint | 1.06 | 1.16 | 216 | 1.01 | 1.02 | 486 | 1.01 | 1.02 | 219 |
| recombs | 1.04 | 1.1 | 254 | 1.01 | 1.03 | 559 | 1.00 | 1.01 | 229 |
| noncompats | 1.02 | 1.04 | 406 | 1.01 | 1.03 | 1290 | 1.01 | 1.04 | 518 |
| arglen | 1.06 | 1.16 | 348 | 1.08 | 1.21 | 459 | 1.05 | 1.12 | 319 |
| Multivariate | 1.15 |  |  | 1.1 |  |  | 1.05 |  |  |

**Table S2.**
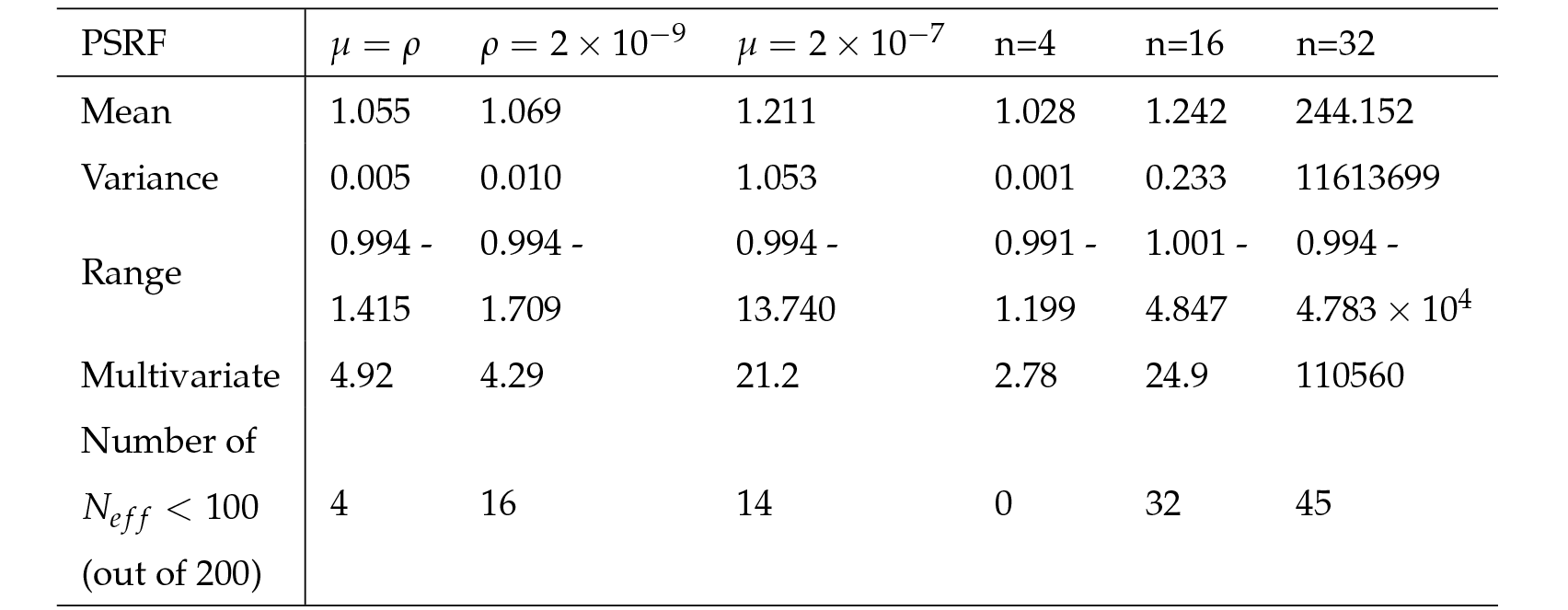
Potential scale reduction factor (PSRF) mean, variance and range for each of 200 coalescence times in ARGweaver, the multivariate PSRF (Plummer et al. 2006) and the number of coalescence times for each the effective sample size (*N*_*e f f*_) is smaller than 100. Unless otherwise noted, mutation rate (*µ*) and recombination rate (*ρ*) are 2 *×* 10^−8^ and sample sizes (n) are 8 haplotypes.

| PSRF | $\mu = \rho$ | $\rho = 2 \times 10^{-9}$ | $\mu = 2 \times 10^{-7}$ | n=4 | n=16 | n=32 |
| --- | --- | --- | --- | --- | --- | --- |
| Mean | 1.055 | 1.069 | 1.211 | 1.028 | 1.242 | 244.152 |
| Variance | 0.005 | 0.010 | 1.053 | 0.001 | 0.233 | 11613699 |
| Range | 0.994 -<br>1.415 | 0.994 -<br>1.709 | 0.994 -<br>13.740 | 0.991 -<br>1.199 | 1.001 -<br>4.847 | 0.994 -<br>$4.783 \times 10^4$ |
| Multivariate | 4.92 | 4.29 | 21.2 | 2.78 | 24.9 | 110560 |
| Number of<br>$N_{eff} < 100$<br>(out of 200) | 4 | 16 | 14 | 0 | 32 | 45 |

**Table S3.**
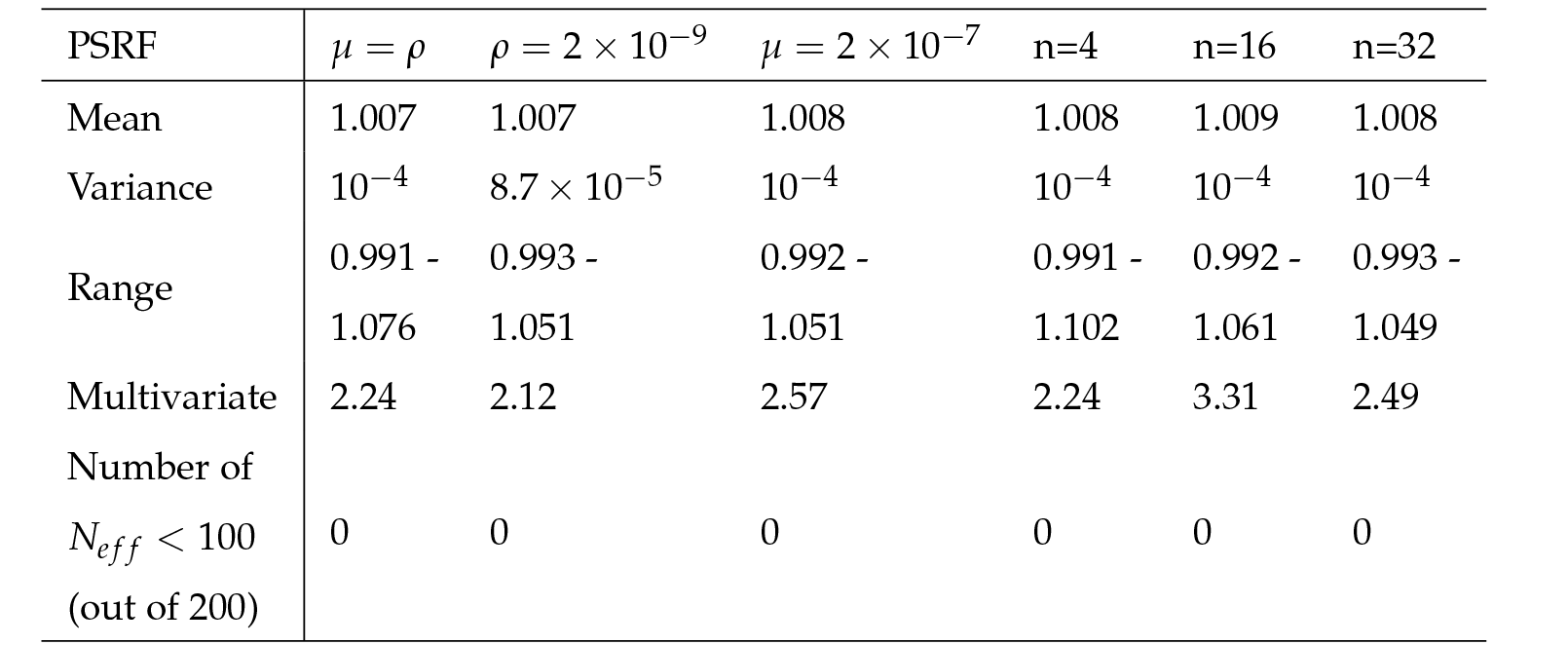
Potential scale reduction factor (PSRF) mean, variance and range for each of 200 coalescence times in Relate, the multivariate PSRF (Plummer et al. 2006) and the number of coalescence times for each the effective sample size (*N*_*e f f*_) is smaller than 100. Unless otherwise noted, mutation rate (*µ*) and recombination rate (*ρ*) are 2 *×* 10^−8^ and sample sizes (n) are 8 haplotypes.

**Table S4.** Minimum and maximum acceptance rates of ARGweaver subtree sampling steps for each simulation.

| Simulation | Acceptance rates |  |
| --- | --- | --- |
|  | Min | Max |
| $n=8; \mu = \rho = 2 \times 10^{-8}$ | 0.283 | 0.532 |
| $n=8; \mu = 2 \times 10^{-8}; \rho = 2 \times 10^{-9}$ | 0.838 | 0.965 |
| $n=8; \mu = 2 \times 10^{-7}; \rho = 2 \times 10^{-8}$ | 0.345 | 0.582 |
| $n=4; \mu = \rho = 2 \times 10^{-8}$ | 0.262 | 0.511 |
| $n=16; \mu = \rho = 2 \times 10^{-8}$ | 0.299 | 0.567 |
| $n=32; \mu = \rho = 2 \times 10^{-8}$ | 0.307 | 0.568 |

**Table S5.** Comparison of ARGweaver results with simulations under infinite sites mutational model and Jukes-Cantor finite sites mutational model, including simulations with values of mutation to recombination rate ratio in between the ones shown in the main text. * indicate results shown in the main text and presented here again for comparison.

| $\mu/\rho$ | Point estimates (MSE) | | Ranks (KLD) | |
| --- | --- | --- | --- | --- |
|  | Infinite sites | Finite sites (JC) | Infinite sites | Finite sites (JC) |
| $\frac{2 \times 10^{-8}}{2 \times 10^{-8}} = 1$ | 0.397* | 0.396 | 0.027* | 0.026 |
| $\frac{4 \times 10^{-8}}{2 \times 10^{-8}} = 2$ | 0.285 | 0.285 | 0.049 | 0.053 |
| $\frac{8 \times 10^{-8}}{2 \times 10^{-8}} = 4$ | 0.195 | 0.197 | 0.113 | 0.112 |
| $\frac{2 \times 10^{-7}}{2 \times 10^{-8}} = 10$ | 0.117* | 0.120 | 0.350* | 0.353 |
| $\frac{2 \times 10^{-8}}{2 \times 10^{-9}} = 10$ | 0.120* | 0.119 | 0.286* | 0.291 |

### Evaluating MCMC Convergence

To evaluate MCMC convergence in ARGweaver and Relate, we run these programs five independent times for the same simulated sequence of 5Mb. We do this for each simulation scenario and evaluate convergence by analysing various statistics extracted at each iteration. For ARGweaver, we analyse statistics from in the .*stats* file, described below. Relate does not generate a similar output, so we extract a subset of the pairwise coalescence times at each MCMC iteration to evaluate convergence. We also evaluate convergence based on selected pairwise coalescence times in ARGweaver, for comparison. Using these statistics extracted at each iteration, we evaluate MCMC convergence by analysing 1) trace plots, 2) autocorrelation plots, 3) effective sample sizes (Taboga 2017; Roy 2020), and 4) potential scale reduction factor (PSRF) (Gelman and Rubin 1992). Analyses and plots were done in R using the function *acf* for autocorrelation, and R package *coda* (Plummer et al. 2006) for effective sample sizes and potential scale reduction factor. These results were used to inform our decisions on burn-in and thinning for MCMC, as well as interpreting results of our evaluations of the methods under different simulated conditions.

#### ARGweaver

##### Convergence of likelihoods

ARGweaver’s *arg-sample* program outputs a .*stats* file containing several statistics for each MCMC iteration: log probability of the sampled ARG given the model (“prior”, in Table S1), log probability of the data given the sampled ARG (“likelihood”), total log probability of the ARG and the data (“joint”), number of recombination events in the sampled ARG (“recombs”), the number of variant sites that cannot be explained by a single mutation under the sampled ARG (“noncompats”), total length of all branches summed across sites (“arglen”) (Hubisz and Siepel 2020). We generated trace plots and calculated autocorrelation between consecutive samples using the likelihood per iteration (Figures S9 and S11). Following visual inspection of these plots, we chose a burn-in consisting of the first 200 samples in most simulations, except in simulations with 10 times higher mutation rate (Figure S9C,F) or sample sizes larger than 8 haplotypes (Figure S11B,C,E,F), where we chose a burn-in of 1200 samples since those chains took longer to converge. In both cases, we ran MCMC for 1000 iterations after burn-in. Based on autocorrelation plots (Figure S9, S11) and on effective sample sizes (Table S1), we thinned ARGweaver samples by recording every 10th MCMC iteration, thus retaining a total of 100 MCMC samples.

Results of the potential scale reduction factor suggested convergence of ARGweaver in simulations with mutation rate equal to recombination rate, with decreased recombination rate and with increased mutation rate (Table S1) - see section below on convergence of individual coalescence times.

##### Convergence of coalescence times

For comparison with Relate, which does not output statistics for each iteration, we also analyse convergence of pairwise coalescence times in ARGweaver. To this end, we extract from each MCMC iteration the values of coalescence times between two pairs of samples at 100 sites equally spaced by 50 kb along the 5Mb simulated sequences. We use those 200 values for convergence diagnostics. Figure S12 shows trace plots of 10 of those sites, for one pair of samples. To evaluate convergence, we calculate potential scale reduction factor (PSRF) for each of the 200 coalescence times, and compare their mean, variance and range (Table S2) among different simulations. In Table S2 we also compare the number of coalescence times that have effective sample sizes lower than 100 (which is our MCMC sample size). These results also lead us to conclude that ARGweaver runs with mutation rate equal to recombination rate have converged. However, in contrast to the results on convergence for statistics recorded in the ARGweaver *stats* files (Table S1), the evaluation of convergence based on coalescence times does not support a conclusion of full convergence for the other simulated data sets. In particular, simulations with mutation to recombination rate ratio of 10 had a large number of coalescence times with effective sample size smaller than 100. The same was true for simulations with 16 and 32 haplotypes. The maximum values of PSRF in those simulations are also further from one, thus indicating a lack of convergence for some coalescence times.

#### Relate

Relate estimates branch lengths using an MCMC algorithm with built in burn-in (Speidel *et al*. (2019) Supplementary Note on Method details 4.2, p. 13). To obtain samples from the posterior distribution, the tree sequence estimated in this first step was used as a starting point. Therefore, we did not implement any extra burn-in to obtain samples from the posterior. Visual inspection of traces plots also suggested that additional burn-in was not necessary (Figure S13).

We evaluated Relate’s MCMC convergence by running it 5 times for each sequence of 5Mb simulated under each set of parameters. We then extracted a subset of pairwise coalescence times to calculate the potential scale reduction factor and effective sample sizes as described above for ARGweaver. We extracted coalescence times for two pairs of samples at 100 equally spaced sites along the sequence (*i*.*e*. separated by 50kb). Table S3 shows these results, which indicate convergence of all Relate runs in all simulated datasets.

### Tsdate prior grid

We ran tsdate with different prior grids, using the function tsdate.build_prior_grid(). The observation that dates inferred by tsdate seem to be bounded to a low maximum value still holds when changing prior grids to have more points (timepoints=100, Figure S14) or when manually specifying time slices with a maximum value of 12 (timepoints=np.geomspace(1e-5, 12, 50), Figure S15).

### ARGweaver subtree sampling acceptance rates

As suggested by ARGweaver authors (Melissa Hubisz and Adam Siepel, personal communication), we have verified that acceptance rates of subtree sampling steps of ARGweaver are within a range that indicates good mixing of the chain, between 10% and 90% (Table S4). All simulations except for the one with reduced recombination rate were within that range. For a visualization of the spread of the values of acceptance rate, Figure S16 shows the acceptance rates for subtree sampling steps of ARGweaver in one 5Mb region of each simulation.

### Additional simulations results for ARGweaver

#### SMC and SMC’ modes in ARGweaver

In all results shown in the main text, we simulated under the standard Hudson (1983) coalescent with recombination, and did inference in ARGweaver under SMC’. Here, we asked whether deviations observed in the posterior distribution of ARGweaver can be explained by differences between the models used for simulation and inference. For this, we simulate sequences in msprime under the SMC and SMC’ models, and run ARGweaver inference using the same model used in the simulation. We simulated 8 haplotypes with mutation rate and recombination rate 2 *×* 10^−8^. Results improve when simulating under SMC’ and inferring under SMC’ (Figures S17B, S18B). Surprisingly, simulating and inferring under SMC (Figures S17A, S18A) is not better than simulating under the full coalescent with recombination model and inferring under SMC (Figures 4, 5).

#### Intermediate values of mutation to recombination rate ratio

Rasmussen *et al*. (2014) mention in their Figure S5 that the quality of ARGweaver estimates generally improved in their simulations with increased mutation to recombination rates ratio (*µ*/*ρ*), but only up to *µ*/*ρ* = 4. Motivated by this observation, we additionally ran simulations with values of *µ*/*ρ* in between the ones shown in the main text (*µ*/*ρ*=1 or *µ*/*ρ*=10), including *µ*/*ρ*=2 and 4. We summarize our results under these conditions in Table S5. We observed a similar pattern for these intermediate values of *µ*/*ρ* = 2, 4 as we had observed from 1 to 10, *i*.*e*. point estimates improve with increased ratio (shown by lower MSE in Table S5), and calibration of the posterior distribution worsens with an increased ratio (show by higher KLD in Table S5).

#### Jukes-Cantor mutational model

In all results shown in the main text, we simulated mutations using an infinite sites model. ARGweaver, on the other hand, uses a Jukes and Cantor (1969) mutational model. Therefore, we hypothesize that differences in the mutational model between simulations and inference could explain deviations in the posterior distribution of ARGweaver, especially in simulations with increased mutation to recombination ratio (*µ*/*ρ*). We found that ARGweaver results with simulations under the Jukes and Cantor (1969) model are very similar to the results under the infinite sites model and follow the same pattern under increased *µ*/*ρ* (Table S5, Figures S20, S21).

## Notes

### Competing Interest Statement

The authors have declared no competing interest.

### Summary of Updates

Supplementary figures' labels and legends updated.

